# Host ecology regulates interspecies recombination in bacteria of the genus *Campylobacter*

**DOI:** 10.1101/2021.08.24.457495

**Authors:** Evangelos Mourkas, Koji Yahara, Sion C. Bayliss, Jessica K. Calland, Håkan Johansson, Leonardos Mageiros, Zilia Y. Muñoz-Ramirez, Grant Futcher, Guillaume Méric, Matthew D. Hitchings, Santiago Sandoval-Motta, Javier Torres, Keith A. Jolley, Martin C. J. Maiden, Patrik Ellström, Jonas Waldenström, Ben Pascoe, Samuel K. Sheppard

**Affiliations:** The Milner Centre for Evolution, Department of Biology and Biochemistry, University of Bath, Bath BA2 7AY, UK; Antimicrobial Resistance Research Center, National Institute of Infectious Diseases, Tokyo, 162-8640, Japan; Centre for Ecology and Evolution in Microbial Model Systems, Linnaeus University, Kalmar, 391 82, Sweden; Unidad de Investigacion en Enfermedades Infecciosas, UMAE Pediatria, Instituto Mexicano del Seguro Social; Swansea University Medical School, Swansea University, Singleton Park, Swansea, SA2 8PP, UK; Department of Zoology, University of Oxford, South Parks Road, Oxford, OX1 3PS, UK; Department of Medical Sciences, Zoonosis Science Centre, Uppsala University, Uppsala, Sweden; Faculty of Veterinary Medicine, Chiang Mai University, Chiang Mai 50100, Thailand

**Author notes:** Corresponding authors: Samuel K. Sheppard. Cambridge Baker Systems Genomics Initiative, Baker Heart and Diabetes Institute, 75 Commercial Rd, Melbourne 3004, Victoria, Australia; Department of Infectious Diseases, Central Clinical School, Monash University, Melbourne, Victoria 3004, Australia (Guillaume Méric).

**Keywords:** *Campylobacter*, genus, species, niche, adaptation, host, evolution

## Abstract

Horizontal gene transfer (HGT) can allow traits that have evolved in one bacterial species to transfer to another. This has potential to rapidly promote new adaptive trajectories such as zoonotic transfer or antimicrobial resistance. However, for this to occur requires gaps to align in barriers to recombination within a given time frame. Chief among these barriers is the physical separation of species with distinct ecologies in separate niches. Within the genus *Campylobacter* there are species with divergent ecologies, from rarely isolated single host specialists to multi-host generalist species that are among the most common global causes of human bacterial gastroenteritis. Here, by characterising these contrasting ecologies, we can quantify HGT among sympatric and allopatric species in natural populations. Analysing recipient and donor population ancestry among genomes from 30 *Campylobacter* species we show that cohabitation in the same host can lead to a 6-fold increase in HGT between species. This accounts for up to 30% of all SNPs within a given species and identifies highly recombinogenic genes with functions including host adaptation and antimicrobial resistance. As described in some animal and plant species, ecological factors are a major evolutionary force for speciation in bacteria and changes to the host landscape can promote partial convergence of distinct species through HGT.

## Introduction

It is well established that bacteria do not conform to a strict clonal model of reproduction but engage in regular horizontal gene transfer (HGT) [1]. This lateral exchange of DNA can confer new functionality on recipient genomes, potentially promoting novel adaptive trajectories such as colonization of a new host or the emergence of pathogenicity [2]. In some cases, gene flow can occur at such magnitude, even between different species [3, 4], that one may question why disparate lineages do not merge and why distinct bacterial species exist at all [5]. An answer to this lies in considering the successive processes that enable genes from one strain to establish in an entirely new genetic background.

The probability of HGT is governed by the interaction of multiple factors, including exposure to DNA, the susceptibility of the recipient genome to DNA uptake, and the impact of recombined DNA on the recipient strain. These factors can be broadly defined in three functional phases and HGT can only occur when gaps align in each successive ecological, mechanistic and adaptive barriers within a given time frame (Figure 1). In the first phase, the quantity of DNA available to recipient strains is determined by ecological factors such as the distribution, prevalence and interactions of donor and recipient bacteria, as well as the capacity for free DNA to be disseminated among species/strains. In the second phase, there are mechanistic barriers to HGT imposed by the homology dependence of recombination [6] or other factors promoting DNA specificity - such as restriction-modification, CRISPR interference or antiphage systems [7–11] - that can act as a defence against the uptake of foreign DNA (mechanistic barriers) [12, 13]. Finally, the effect that HGT has on the fitness of the recipient cell in a given selective environment (adaptive barrier) will determine if the recombinant genotype survives for subsequent generations [2, 14].

**Figure 1.**
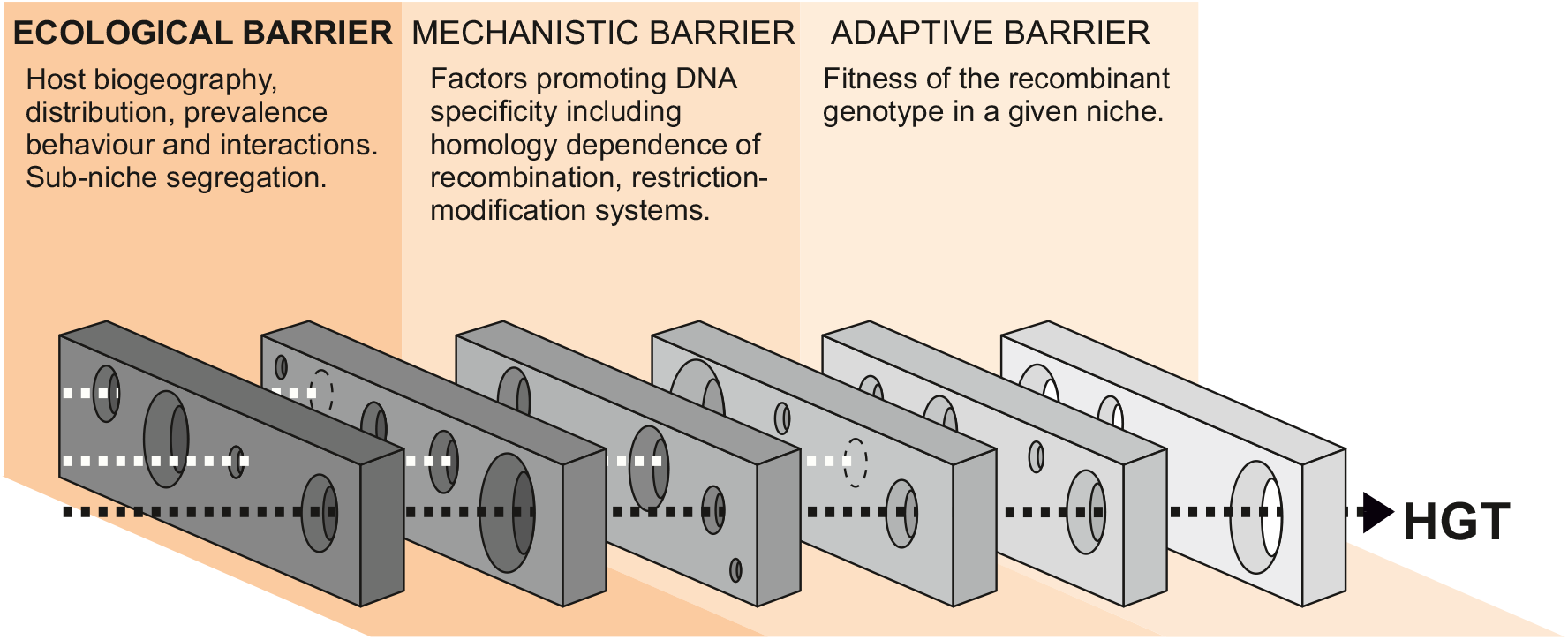
Barriers to horizontal gene transfer in bacteria. A series of barriers must be surmounted for DNA to transmit from one species to another. These are broadly defined in three categories. At a given time, alignment of holes in successive barriers is necessary for HGT to occur. Here we focus on ecological barriers that are influenced by multiple factors that reflect the physical isolation of bacteria in separate niches.

Understanding how ecology maintains, and potentially confines, distinct strains and species has become increasingly important in the light of global challenges such as the emergence and spread of zoonotic pathogens [15]. A typical approach to investigating this is to consider spillover of particular strains or clones from one host to another (clonal transmission). This is an important phenomenon and may be influenced by anthropogenic change, such as habitat encroachment or agricultural intensification [16]. However, in many cases, important phenotypes, including antimicrobial resistance (AMR) [17–19], can be conferred by relatively few genes. In such cases, it may be important to consider how cohabiting strains and species can potentially draw genes from a common pangenome pool [20–23] and how genes, rather than clones, can transition between segregated populations (gene pool transmission). To investigate the impact of ecological segregation (ecological barriers) on this gene pool transmission, in natural populations, requires quantification of HGT among sympatric and allopatric bacteria.

Species within the genus *Campylobacter* are an ideal subject for considering how ecology influences the maintenance of genetically distinct species for several reasons. First, *Campylobacter* are a common component of the commensal gut microbiota of reptiles [24, 25], birds [26, 27] and mammals [28] but, being microaerophilic, do not survive well outside of the host. This creates island populations that have some degree of ecological isolation. Second, because at least 12 species have been identified as human pathogens [29] and *C. jejuni* and *C. coli* among the most common global causes of bacterial gastroenteritis [30], large numbers of isolate genomes have been sequenced from potential reservoir hosts as part of public health source tracking programs [31, 32]. Third, within the genus there are species and strains that inhabit one or multiple hosts (ecological specialists and generalists [16, 26, 33–37]). As a single host can simultaneously carry multiple lineages [38], possibly occupying different sub-niches within that host [39], there is potential to compare allopatric and sympatric populations. Finally, high magnitude interspecies admixture (introgression) between *C. jejuni* and *C. coli* isolated from agricultural animals suggests that host ecology plays a role in the maintenance of species [40–43].

Here, we quantify HGT among 600 genomes from 30 *Campylobacter* species using a ‘chromosome painting’ approach [44–46] to characterize shared ancestry among donor and recipient populations. Specifically, we investigate the role of ecological barriers to interspecies gene flow. By identifying recombining species pairs within the same and different hosts we can describe interactions where co-localization enhances gene flow, quantify the impact of ecological barriers in these populations and distinguish highly recombinogenic genes that are found in multiple genetic backgrounds. This provides information about the evolutionary forces that gave rise to species and the extent to which ecological barriers maintain them as discrete entities.

## Results

### Host restricted and host generalist *Campylobacter* species

Isolate genomes were taken from publicly available databases to represent diversity within the genus *Campylobacter,* including environmental isolates from the closely related *Arcobacter* and *Sulfurospirillum* species to provide phylogenetic context within the *Campylobacteraceae* family (Figure 2a-figure supplement 1). In total, there were 631 isolates from 30 different *Campylobacter* species (Figure 2a) and 64 different sources, isolated from 31 different countries between 1964 and 2016 (Supplementary File 1). Among the isolates, 361 were *C. jejuni* and *C. coli* and could be classified according to 31 Clonal Complexes (CCs) based upon sharing four or more alleles at seven housekeeping genes defined by multi-locus sequence typing (MLST) (Supplementary File 1) [47] and were representative of known diversity in both species [16, 33]. The obligate human commensal and pathogen *C. concisus* (n=106 isolates), comprised 2 genomospecies (GSI, n=32 and GSII, n=74), as previously described [48] (Supplementary File 1). The collection also included 52 *C. fetus* isolate genomes, including 3 subspecies: *C. fetus subsp. fetus* (n=8), *C. fetus subsp. venerealis* (n=23) and *C. fetus subsps. testudinum* (n=21) (Supplementary File 1) [49]. Two clades were observed in *C. lari* (Figure 2a-figure supplement 2) which could correspond to previously described subspecies based on 16S rRNA sequencing [50].

**Figure 2.**
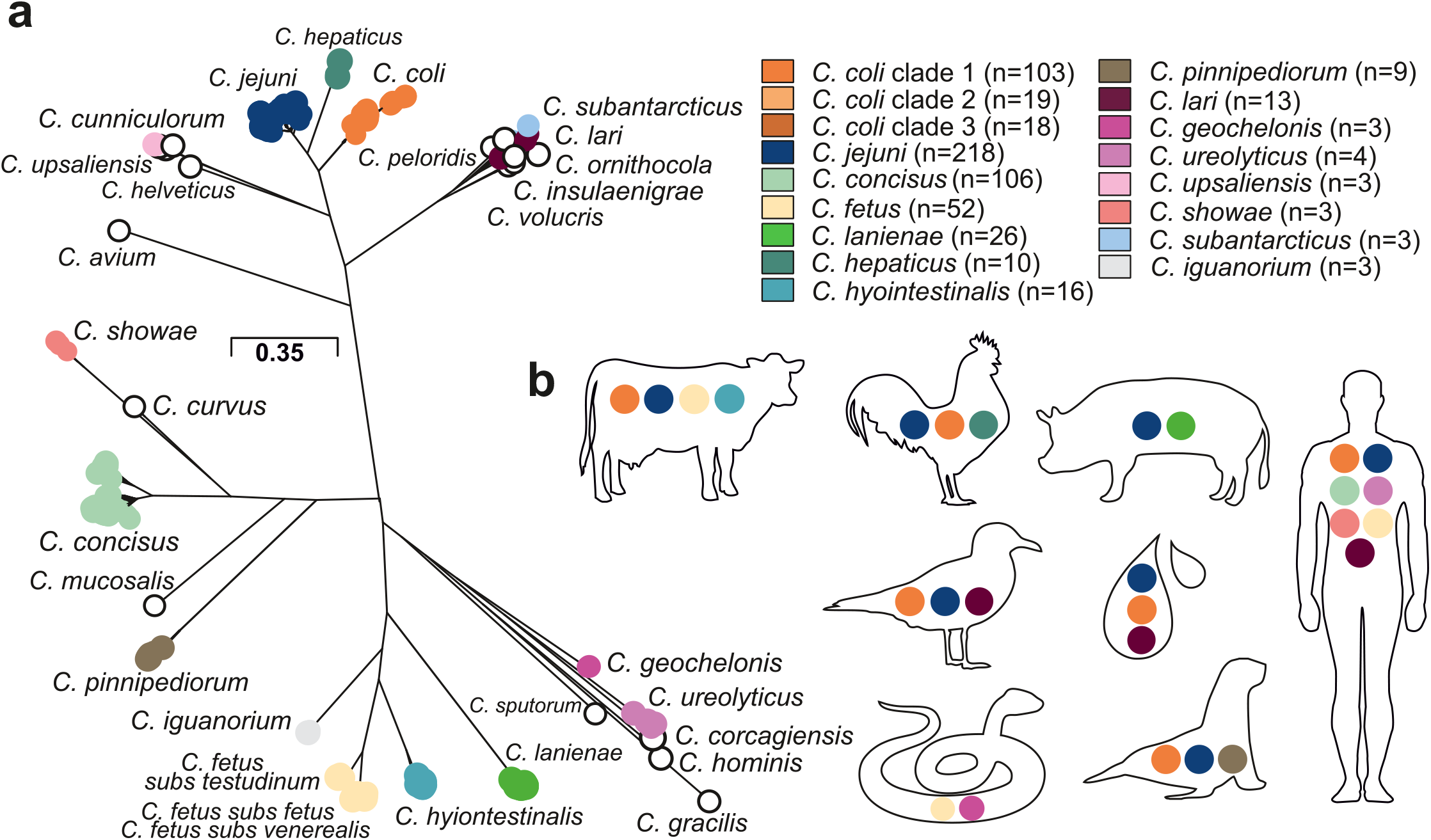
Population structure and host ecology in the genus *Campylobacter*. **a**, Phylogenetic tree of 631 *Campylobacter* isolates from 30 species reconstructed using a gene-by-gene concatenated alignment of 820 core genes (shared by >95% of isolates) and an approximation of the maximum-likelihood algorithm (ML) implemented in RAxML. The species name is indicated adjacent to the associated sequence cluster. The scale bar indicates the estimated number of substitutions per site. **b**, Isolation source of *Campylobacter* species with n≥3 isolates.

A maximum-likelihood phylogeny of the *Campylobacter* genus was reconstructed on a gene-by-gene concatenated sequence alignment of 820 gene families shared by >95% of all isolates, with a core genome of 903,753 base pairs (Figure 2a). The phylogeny included species which appear to be restricted to one host or environment, including *C. iguanorium* [51] and *C. geochelonis* [52] (reptiles), *C. lanienae* [53] (pigs), *C. hepaticus* [54] (chicken liver), *C. lari* group [55] (marine birds and environment) and *C. pinnipediorum* [56] (seals) species, most of which were discovered recently (Figure 2a–figure supplement 3). There was no evidence that phylogeography was reflected in the observed population structure for *Campylobacter* isolates from multiple hosts and countries. (Figure 2–figure supplement 4). This is unsurprising as it is well known that host associated genetic variation transcends phylogeographic structuring in *Campylobacter* [35]. While some low-level local gene flow can be identified within a given country [57], this is vastly outweighed by recombination within particular host niches [36], particularly in small isolate collections such as those for some of the species in this study.

Host restricted species had lower diversity possibly linked to low sample numbers, with *C. hepaticus* having the lowest diversity (Figure 2–figure supplement 2) with 8/10 genomes associated with isolates from the same outbreak [54]. For other species there was evidence of a broad host range (ecological generalists) (Figure 2b). For example, highly structured *C. jejuni* and *C. coli* isolates were sampled from seven and six host sources respectively (Figure 2–figure supplements 2-3, Supplementary File 1). For *C. fetus* there was distinct separation between mammal-associated *C. fetus* subsp *fetus* and *C. fetus* subsp *venerealis* and reptile-associated *C. fetus* subsp *testudinum* (Figure 2–figure supplement 2) as previously described [49]. Unsurprisingly, a large proportion of the isolates in this study were from humans, likely reflecting intensive sampling. *C. jejuni* (27.52%; n=60/218), *C. coli* (14.68%; n=32/218) and *C. concisus* (44.5%; n=97/218) were all common among human clinical samples. However, less common species were also present, with nearly half of all *Campylobacter* species (44.83%, n=13/29) isolated from humans at least once (Figure 2b, Supplementary File 1). Agricultural animals were also a common source accounting for more than 1/3 of the isolates (38.35%; 242/631), with 10/30 *Campylobacter* species isolated from more than one source (Figure 2b, Supplementary File 1).

### Evidence of interspecies recombination in the core and accessory genome

Genome size varied between 1.40 and 2.51 Mb (Figure 3–figure supplement 1) (mean 1.73) and the number of genes (per isolate) ranged from 1,293 to 2,170 (mean 1,675) (Figure 3a– figure supplement 2). The pangenome for the genus comprised 15,649 unique genes, found in at least one of the 631 isolates (Figure 3b–source data 1), with 820 genes (5.24 % of the pangenome) shared by >95% of all isolates (core genome), across 30 species (Figure 3b– source data 1). We excluded species with fewer than 3 isolates in subsequent analysis. For the remaining 15 species the core genome ranged in size from 1,116 genes in *C. lari* to 1,700 in *C. geochelonis* (Figure 3a right panel–source data 1). Differences were also noted in the size of accessory genomes, with *C. concisus* (mean: 981 genes)*, C. hyointestinalis* (mean: 946 genes), *C. showae* (mean: 1,160 genes), *C. geochelonis* (mean: 1,021 genes) and *C. fetus* (mean: 912 genes) containing the highest average number of accessory genes (Figure 3a left panel–source data 1). Functional annotation of all 14,829 accessory genes showed that 71% (10,561) encoded hypothetical proteins of unknown function due to the lack of homology with well-characterized genes (Figure 3–figure supplement 3) [58]. Remaining genes were related to metabolism, DNA modification, transporters, virulence, inner membrane/periplasmic, adhesion, regulators, metal transport and antimicrobial resistance (Figure 3–figure supplement 3).

**Figure 3.**
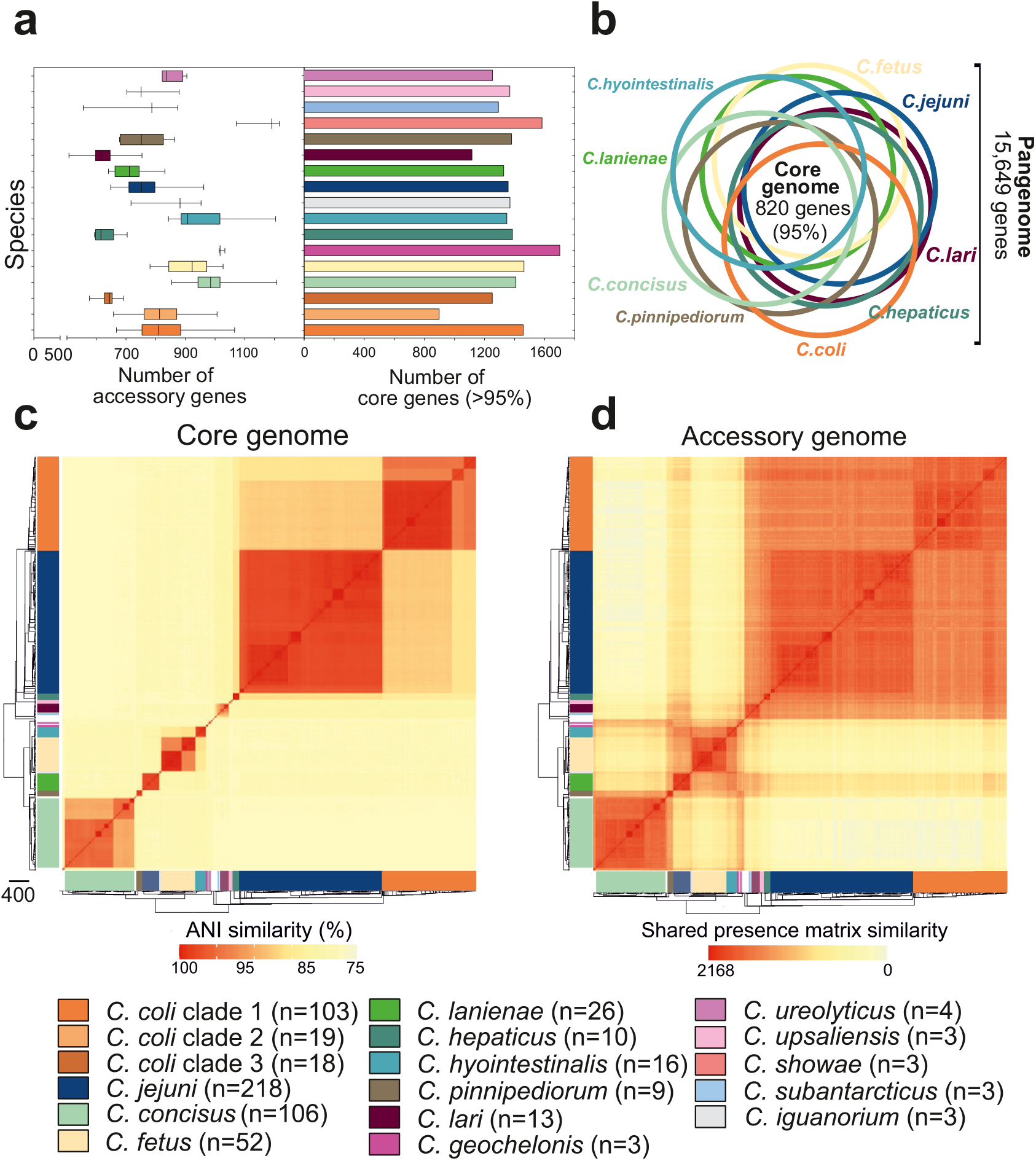
Core and accessory genome variation in the genus *Campylobacter*. **a**, Overall distribution of the total number of accessory genes (left) and core genes (right) per isolate for each *Campylobacter* species (where n≥3 isolates). The number of accessory genes is shown as boxplots (min to max). **b**, Venn diagram of pangenomes among different *Campylobacter* species (n≥9). The number of core genes shared by all species is illustrated in the center. **c**, Pairwise average nucleotide identity comparison calculated for all 631 *Campylobacter* isolates based upon 820 core genes shared by >95% of isolates. ANI values <75% are not calculated by FastANI [59]. **d**, Pairwise accessory genome similarity based upon gene presence or absence at 2,168 non-core loci. The heatmaps coloring ranges from yellow (minimum) to red (maximum). The matrices are ordered according to the phylogenetic tree presented in figure 2a. Different colours correspond to *Campylobacter* species with ≥3 isolates.

To further understand genetic differentiation within and between species, we generated genus-wide similarity matrices for the core and accessory genomes (Figure 3c-d–source data 1). For the core genome, pairwise average nucleotide identity (ANI) was calculated for shared genes in all possible genome pairs (Figure 3c–source data 1) using FastANI [59]. On average, isolates of the same species shared >95% similarity (Figure 3c–source data 1), with decreasing genetic similarity (between 85% and 90%) over greater phylogenetic distances. The number of core genome SNPs ranged from 983 to 230,264 for the 15 *Campylobacter* species with ≥ 3 isolates in our dataset, with *C. coli* and *C. concisus* having the greatest mean SNP numbers (Figure 3–figure supplement 4a) indicating considerable diversity within these species. In contrast *C. hepaticus* and *C. geochelonis* had low mean SNP numbers with 986 and 4,310, respectively. This is likely related to low sample numbers with isolates either sampled in close proximity [52] or from a single outbreak [54].

The core genome similarity matrix provided initial evidence of interspecies gene flow (introgression). This can be observed as elevated nucleotide identity between *C. jejuni* and clade 1 *C. coli* (Figure 3c–source data 1), consistent with previous studies [40, 42, 43]. Further evidence of introgression came from pairwise ANI comparison of genus-wide core genes, in all isolates of the 15 major *Campylobacter* species, to the *C. jejuni* genome (Figure 3–figure supplement 4b). In the absence of gene flow, isolates from the two species should have an approximately unimodal ANI distribution reflecting accumulation of mutations throughout the genome. This was largely the case but for some species, low nucleotide divergence suggested recent recombination with *C. jejuni*. There was also evidence of interspecies accessory genome recombination. Presence/absence patterns in the accessory genome matrix show considerable accessory gene sharing among several species that was inconsistent with the phylogeny (Figure 3d–source data 1). This is well illustrated in *C. lanienae* where much of the accessory genome was shared with other *Campylobacter* species (Figure 3d–source data 1).

### Enhanced interspecies recombination among cohabiting species

For *Campylobacter* inhabiting different host species there is a physical barrier to HGT. However, when there is niche overlap, interspecies recombination can occur, for example between *C. jejuni* and *C. coli* inhabiting livestock [33, 40, 42]. To understand the extent to which inhabiting different hosts impedes interspecies gene flow we quantified recombination among *Campylobacter* species where isolates originated from same host (*x_1_, y*) and different hosts (*x_2_, y*) (Figure 4a).

**Figure 4.**
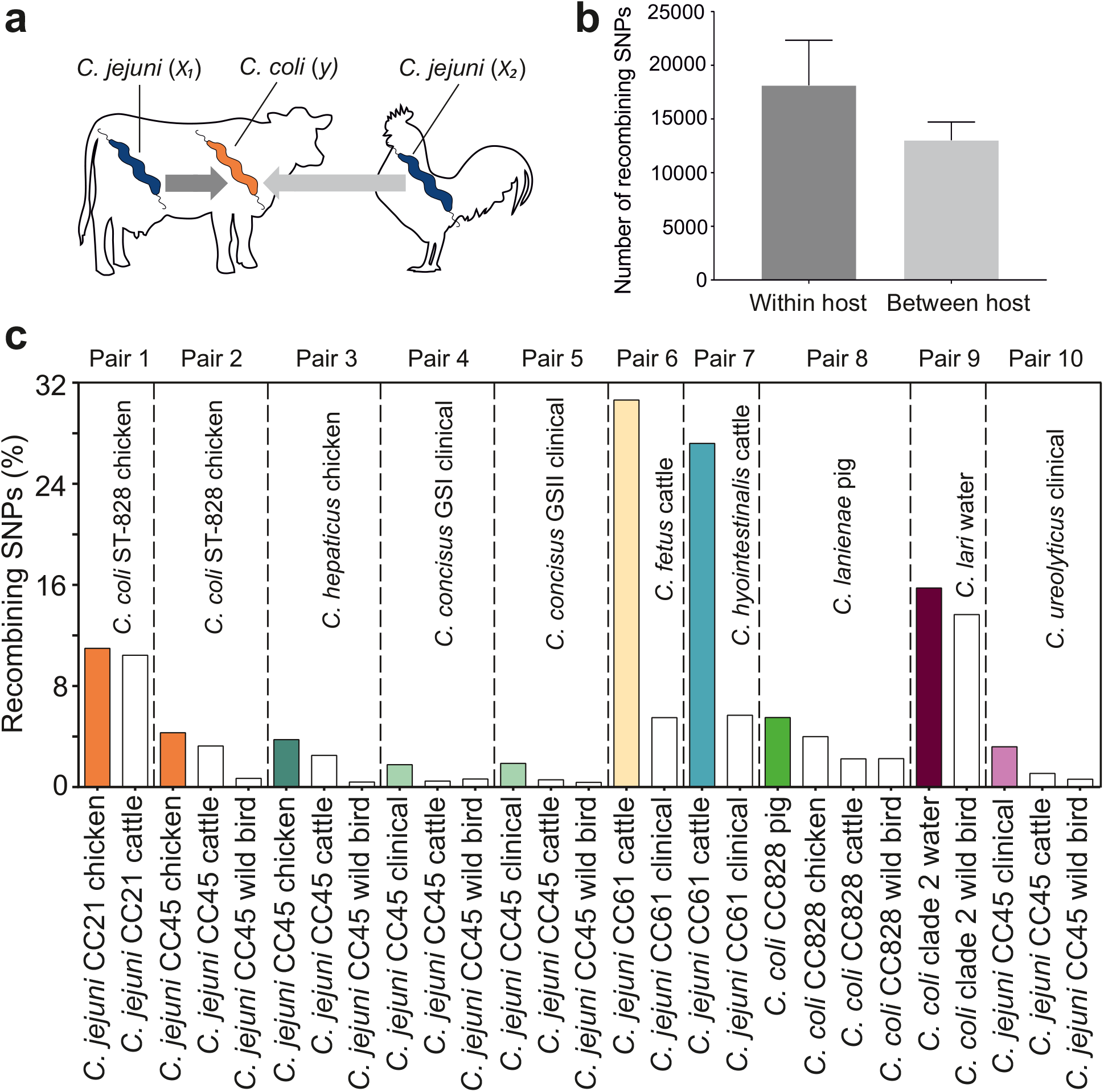
Elevated within-host interspecies recombination and donor-recipient comparisons. **a**, A hypothesis depicting the relationships between *Campylobacter* species, *C. jejuni (x_1_, x_2_)* and *C. coli* (*y*), when found in the same or in different hosts. **b**, Number of recombining SNPs within and between host as inferred by chromosome painting analysis for all donor recipient species comparisons. The error bar represents the standard error of the mean (SEM). **c**, The figure shows the number of donated SNPs in 10 donor-recipient pair species comparisons. The proportion (%) of recombining SNPs with >90% probability of copying from a donor to a recipient genome is illustrated in the *y* axis. All donor groups are shows in the *x* axis. All coloured boxes correspond to comparison where donors and recipients are found in the same host.

ChromoPainterV2 software was used to infer tracts of DNA donated from multiple donor groups, belonging to the same CC but isolated from different hosts to recipient groups (Materials and Methods). Among 27 combinations of multiple donor groups and recipient groups, overall, there were more recombining SNPs within hosts than between hosts (Figure 4b) and for 10/27 species pairs there was evidence of enhanced within species recombination (*x_1_→y* > *x_2_→y*; Figure 4c). To assess the robustness of the analysis we included the effect of randomization and repeated the analysis by assigning random hosts for every strain (Figure 4–figure supplement 1). In the 10 pair species comparisons where *x_1_→y* > *x_2_→y*, we detected 174,594 within-host recombining SNPs (mapped to 473 genes; 28.8% of NCTC11168 genes) and 109,564 between-host recombining SNPs (mapped to 395 genes; 24.05% of NCTC11168 genes). From the 473 within-host recombining genes, 45 genes contained the highest number (>95^th^ percentile) of recombining SNPs (Figure 4–figure supplements 2–3, Supplementary File 2). These genes have diverse inferred functions including metabolism, cell wall biogenesis, DNA modification, transcription, and translation (Supplementary File 2).

Interspecies recombination was observed for isolates sampled from chickens between generalist lineages CC21 and CC45 (donors; *C. jejuni*) and generalist CC828 (recipient; *C. coli*). These lineages appear to have high recombination to mutation (*r/m*) ratio as inferred by ClonalFrameML (Supplementary File 3). DNA from generalist *C. jejuni* CC45 was introduced into three *Campylobacter* species, including *C. hepaticus* (chicken), *C. concisus* GSI and GSII (clinical) and *C. ureolyticus* (clinical) (Figure 4c–figure supplement 2–3, Supplementary File 4). Clonal complex 45 had the highest *r/m* ratio from all other lineages or species involved in the comparisons (Supplementary File 3). There was increased recombination in genomes sampled from cattle between *C. jejuni* CC61 (donor; *C. jejuni*) and *C. fetus* and *C. hyointestinalis* (recipients) with 71.75% of all within-host recombining SNPs from all 10 comparisons detected in these two pairs (Figure 4c–figure supplement 2–3, Supplementary File 4). Agricultural associated *C. jejuni* CC61 and *C. fetus* subsp. *venerealis* involved in these comparisons were among the lineages and subspecies with the highest *r/m* ratios (Supplementary File 3). The cattle-associated CC61 has previously been described as highly recombinant, and has been associated with rapid clonal expansion and adaptation in cattle [16].

### The within-host mobilome

Bacteria inhabiting the same niche may benefit from functionality conferred by similar gene combinations. Recombination can promote the dissemination of adaptive genetic elements among different bacterial species. Therefore, we postulated that the genes that recombine most among species (>95^th^ percentile) will include those that are potentially beneficial in multiple genetic backgrounds. To investigate this, we quantified mobility within the genome identifying recombining SNPs found in more than one species comparison (Figure 5a). These SNPs mapped to 337 genes (20.52% of the NCTC11168 genes; 2.15% of the pangenome) (Figure 5a, Supplementary File 5). We found that 32 of those genes (9.49%) have also been found on plasmids (Supplementary File 5). A total of 16 genes showed elevated within-host interspecies recombination in more than five species pairs (Figure 5c, Supplementary File 5). Genes included *cmeA* and *cmeB* which are part of the predominant efflux pump CmeABC system in *Campylobacter*. Sequence variation in the drug-binding pocket of the *cmeB* gene has been linked to increased efflux function leading to resistance to multiple drugs [60]. Many of the same antimicrobial classes are used in human and veterinary medicine and this may be linked to selection for AMR *Campylobacter*, that are commonly isolated from livestock [61]. To investigate this further, we compared the genomes of all 631 isolates in our dataset to 8,762 known antibiotic resistance genes from the Comprehensive Antibiotic Resistance Database (CARD) [62], ResFinder [63] and the National Center for Biotechnology Information (NCBI) databases. Homology (>75%) was found for 42 AMR determinants associated with multi-drug efflux pumps, aminoglycosides, tetracyclines and β-lactams (Figure 5b–figure supplement 1–source data 1). Species that contained >40% isolates from livestock, including *C. jejuni, C. coli, C. lanienae, C. hepaticus, C. hyointestinalis* and *C. fetus* contained far more AMR determinants (Figure 5d–figure supplement 1–source data 1). AMR genes are often collocated in the genome [64] and our analysis revealed several gene clusters (Figure 5–figure supplement 2) that have been described in previous studies [64, 65]. These findings are consistent with HGT-mediated circulation of AMR genes among different *Campylobacter* species and support hypotheses that ecology drives gene pool transmission [2, 64].

**Figure 5.**
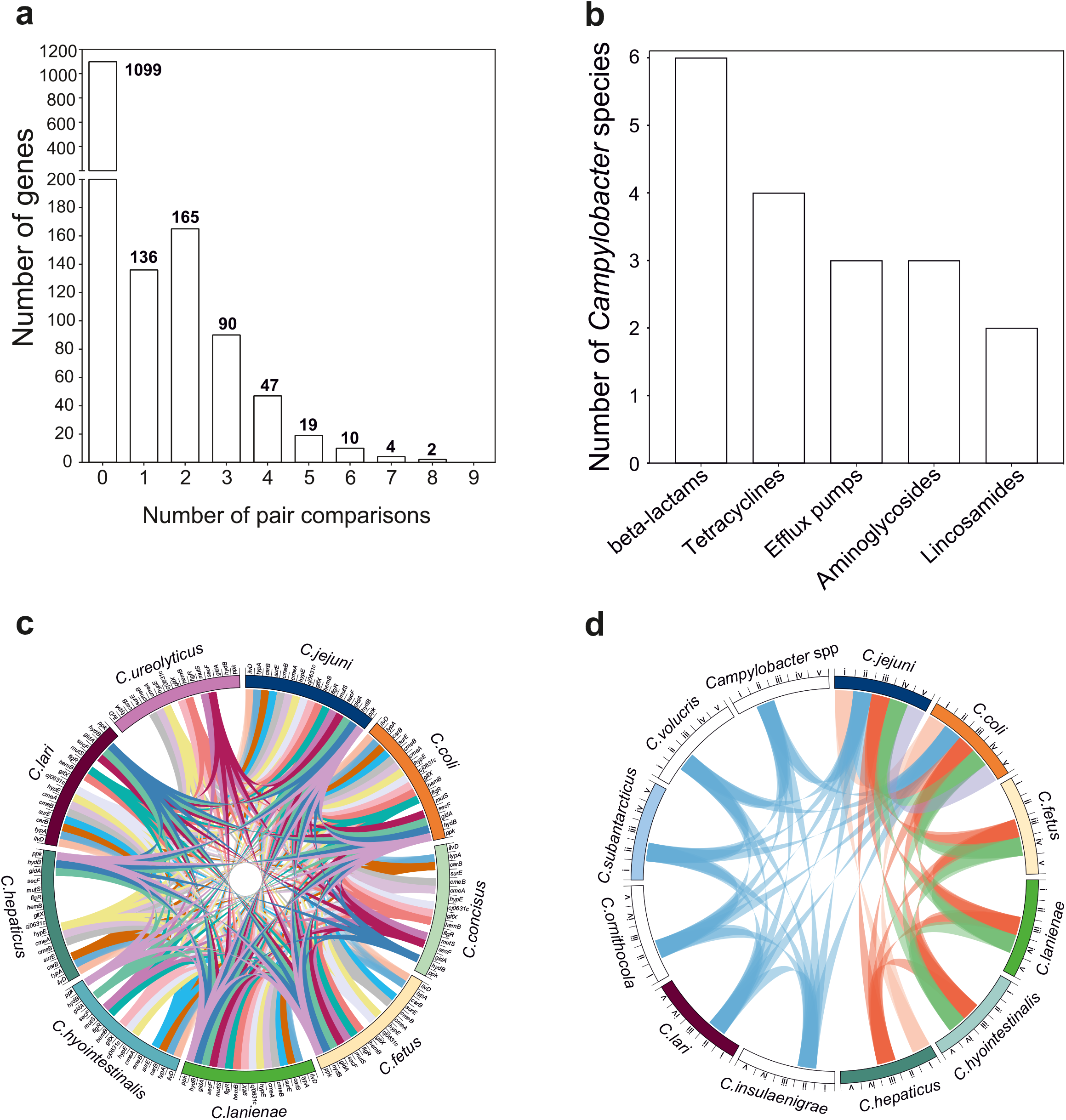
The mobilome of the *Campylobacter* genus. **a**, The graph illustrates the proportion of recombining genes in 10 different species comparisons. The number of species pairs in which the gene was found to recombine is shown on the *x* axis and the number of genes in each category is given on the *y* axis and. The exact number of genes found in each group comparison is shown on the top of each box. **b**, Number of *Campylobacter* species harbouring AMR genes that belong to efflux pumps and four different antibiotic classes which are shown on the *x* axis. **c**, The circos plot indicates the 16 genes involved in recombination in >5 donor-recipient pair species comparisons. Gene matches are indicated by joining lines, coloured differently for each gene. Gene names are shown around the perimeter for each *Campylobacter* species. **d**, The circos plot indicates the sharing of AMR genes associated with efflux pumps and four antibiotic classes among *Campylobacter* species. Presence of at least one gene (not necessarily the same gene) conferring resistance to a specific antibiotic class is indicated by joining lines, coloured differently for each drug class. Efflux pumps (i), β-lactams (ii), tetracyclines (iii), aminoglycosides (iv) and lincosamides (v) are shown around the perimeter for each *Campylobacter* species.

*Campylobacter* host transmission and virulence have been linked with biofilm formation and changes into surface polysaccharides [66, 67]. The *carB* gene showed elevated within-host interspecies recombination in eight species pair comparisons (Figure 5c, Supplementary File 5). This gene encodes a carbamoylphosphate synthase that has been associated with biosynthesis of substrates for many polysaccharides and is known to contain transposon insertion sites upstream of its genomic position [67]. Other genes with elevated within-host interspecies transfer (>7 species pairs) included *typA* (Figure 5c, Supplementary File 5), a translator regulator for GTPase and *gltX* (Figure 5c), *a glutamate-tRNA ligase*, promoting survival under stress conditions [68, 69]. Other genes included *gidA* and *hydB* associated with virulence [70] and hydrogenase enzyme activity (respiratory pathway in *C. concisus,* 69), respectively. By considering genes that overcome barriers to interspecies recombination and establish in multiple new genetic backgrounds, it may be possible to infer important phenotypes that allow bacteria to adapt to different hosts and environments.

## Discussion

Phylogenetic reconstruction of the genus *Campylobacter* revealed a highly structured population. Distinct core genome clustering largely supported known classification for species, subspecies (*C. fetus*, [49]), genomospecies (*C. concisus*, [48]) and clades (*C. coli* [42]). Also consistent with previous studies, certain species are principally associated with a specific host niche. For example, *C. fetus subsp testudinum*, *C. iguanorium, C. geochelonis* were only sampled from reptile species, and *C. pinnipediorum* was only sampled from seals. However, for several species there was clear evidence for host generalism, including *C. jejuni, C. coli* and *C. lari*, all of which were sampled from multiple hosts [26, 72]. It is clear that the hosts with the greatest diversity of *Campylobacter* species were agricultural animals (and humans) (Figure 2–figure supplement 3). While this undoubtedly reflects oversampling of these sources to some extent, the cohabitation of species in the same host niche potentially provides opportunities for interspecies HGT.

Initial evidence of interspecies gene flow came from comparison of average nucleotide identity (ANI) and the accessory genome gene presence/absence for all isolates. In each case, patterns of genetic similarity largely mirrored the phylogeny. However, consistent with previous studies [40], there was clear evidence of elevated homologous and non-homologous recombination between some species. For example, core genome ANI was higher between *C. jejuni* and *C. coli* clade 1, compared to other *C. coli* clades (Figure 3c–source data 1). The evidence for non-homologous gene sharing was even more striking with accessory genome sharing across considerable genetic distances (Figure 3d–source data 1), exemplified by *C. lanienae* which shares accessory genes with most other *Campylobacter* species.

To quantify the extent to which ecological barriers influenced interspecies gene flow, it was necessary to focus on donor-recipient species pairs where there was evidence of elevated HGT in the same (sympatry) compared to different (allopatry) hosts. Perhaps unsurprisingly, this was not the case for all species comparisons. Interacting factors could lead to genetic isolation even when species inhabit the same host. First, rather than being a single niche, the host represents a collection of subniches with varying degrees of differentiation. For example, gut-associated bacteria in the same intestinal tract have been shown to occupy different microniches [73] and more striking segregation may be expected between *C. hepaticus* inhabits the liver in poultry [54] and gut-dwelling *C. jejuni* and *C. coli* in the same host. Second, there is potential for the resident microbiota to influence the colonization potential of different *Campylobacter* species and therefore the opportunity for genetic exchange, for example through succession [74] and inhibition of transient species by residents, as seen in some other bacteria [75–77] in humans.

Continued exposition of the microecology of subniches is important but for 10 species comparisons there was clear evidence of enhanced within-host gene flow allowing quantitative analysis of ecological barriers to gene flow. Specifically, there was on average a 3-fold increase in recombination among species pairs inhabiting the same host. In some cases, this was greater, with 5-6 times more recombination among cohabiting species *C. jejuni* and *C. hyointestinalis/C. fetus* in cattle. In absolute terms, this equates to approximately 30% of all recorded SNPs in the recipient species being the result of introgression. To place this in context, if greater than half (51%) of the recorded SNPs resulted from interspecies recombination then the forces of species convergence would be greater than those that maintain distinct species. If maintained over time, these relative rates could lead to progressive genetic convergence unless countered by strong genome-wide natural selection against introgressed DNA.

Quantitative SNP-based comparisons clearly ignore one very important factor. Specifically, that recombined genes that do not reduce the fitness of the recipient genome (provide an adaptive advantage) will remain in the population while others will be purged through natural selection. Therefore, by identifying genomic hotspots of recombination and the putative function of genes that recombine between species it is possible to understand more about micro-niche segregation and the host adapted gene pool. Of the 35 genes with evidence of enhanced within host HGT in ≥5 species pairs, several were linked to functions associated with proliferation in, and exploitation of, the host. For example, the *carB* gene, encoding the large subunit of carbamoylphosphatase associated with polysaccharide biosynthesis, recombined in eight cohabiting species pairs and is potentially linked to enhanced virulence and growth [67]. In addition, other highly mobile genes, including *typA* and *gltX* are associated with survival and proliferation in stress conditions [68, 69], and *hydB* is linked to NiFe hydrogenase and nickel uptake that is essential for the survival of *C. jejuni* in the gut of birds and mammals [78].

Some genes showed evidence of elevated recombination in a specific host species. For example, the *glmS* and *napA* genes in cohabiting *Campylobacter* species in cattle. In many bacteria, analogs of *glmS* have multiple downstream integration specific sites (Tn7) [79] which may explain the mobility of this gene. Explaining the mobility of *napA* is less straight forward, but this gene is known to encode a nitrate reductase in *Campylobacter* [80] in microaerobic conditions which may be ecologically significant as the accumulation of nitrate in slurry, straw and drainage water can be potentially toxic to livestock mammals [81].

Factors such as host physiology, diet, and metabolism undoubtedly impose selection pressures upon resident bacteria and the horizontal acquisition of genes provides a possible vehicle for adaptation. However, the widespread use of antimicrobials by humans, pets and livestock production [82, 83], provides another major ecological barrier to niche colonization. We found that *gyrA* was among the most recombinogenic genes in *Campylobacter* in chickens. This is important as a single mutation in this gene is known to confer resistance to ciprofloxacin [84]. While the rising trend in fluoroquinolones resistance in *Campylobacter* from humans and livestock [85] may result from spontaneous independent mutations, it is likely that it is accelerated by HGT. However, there is currently no clear evidence for the transfer of resistant versions of *gyrA.* Interspecies recombination of AMR genes has been observed between *C. jejuni* and *C. coli* isolates from multiple sources including livestock, human and sewage [64]. Consistent with this, we found AMR genes present in strains from 12 *Campylobacter* species in multiple hosts (Figure 5–figure supplement 2). In some cases, strains from phylogenetically closely related species (*C. fetus* and *C. hyointestinalis*) isolated from cattle, shared the same AMR gene cluster (*tet44* and *ant(6)-Ib*) described before in *C. fetus* subsp. *fetus* [65], indicating the circulation of colocalized AMR genes among related species and host niche gene pools. Strikingly, the efflux pump genes *cmeA* and *cmeB*, associated with multidrug resistance (MDR) were highly mobile among *Campylobacter* species with evidence of elevated within host interspecies recombination in >7 species pairs. Furthermore, the *gltX* gene, which when phosphorylated by protein kinases promotes MDR [69], was also among the most introgressed genes. While a deeper understanding of gene interactions, epistasis and epigenetics would be needed to prove that the lateral acquisition of AMR genes promotes niche adaptation, these data do suggest that HGT may facilitate colonization of antimicrobial-rich host environments, potentially favouring the spread of genes into multiple genetic backgrounds.

In conclusion, we show that species within the genus *Campylobacter* include those that are host restricted as well as host generalists. When species cohabit in the same host, ecological barriers to recombination can be perforated leading to considerable introgression between species. While the magnitude of introgession varies, potentially reflecting microniche structure with the host, there is clear evidence that ecology is important in maintaining genetically distinct species. This parallels evolution in some interbreeding eukaryotes, such as Darwin’s Finches, where fluctuating environmental conditions can change the selection pressures acting on species inhabiting distinct niches, potentially favouring hybrids [86, 87]. Consistent with this, the host landscape is changing for *Campylobacter*, with intensively reared livestock now constituting 60-70% of bird and mammal biomass on earth respectively [88]. This creates opportunities for species to be brought together in new adaptive landscapes and for genes to be tested from multiple genetic backgrounds. By understanding the ecology of niche segregation and the genetics of bacterial adaptation we can hope to improve strategies and interventions to reduce the risk of zoonotic transmission and the spread of problematic genes among species.

## Materials & Methods

### Isolate genomes

A total of 631 *Campylobacter*, 17 *Arcobacter*, seven *Sulfurospirillum* and five *Helicobacter* genomes were assembled from previously published datasets (Supplementary File 1). Isolates were sampled from clinical cases of campylobacteriosis and faeces of chickens, ruminants, wild birds, wild mammals, pets and environmental sources. Genomes and related metadata were uploaded and archived in the BIGS database [89]. Quality control was performed based on the genome size, number of contigs, N50 and N95 contig length using the integrated tools in BIGS database. All assembled contigs were further screened for contamination and completeness using CheckM [90] (Supplementary File 1). All assembled genomes can be downloaded from FigShare (doi: 10.6084/m9.figshare.15061017). Comparative genomics analyses focused on the *Campylobacter* genomes representing 30 species including: *C. avium* (n=1); *C. coli* (n=143); *C. concisus* (n=106); *C. corcagiensis* (n=1); *C. cuniculorum* (n=2); *C. curvus* (n=2); *C. fetus* (n=52); *C. geochelonis* (n=3); *C. gracilis* (n=2); *C. helveticus* (n=1); *C. hepaticus* (n=10); *C. hominis* (n=1); *C. hyointestinalis* (n=16); *C. iguanorium* (n=3); *C. insulaenigrae* (n=1); *C. jejuni* (n=218); *C. lanienae* (n=26); *C. lari* (n=13); *C. mucosalis* (n=1); *C. ornithocola* (n=1); *C. peloridis* (n=1); *C. pinnipediorum* (n=9); *C. rectus* (n=1); *C. showae* (n=3); *C. sputorum* (n=1); *C. subantarcticus* (n=3); *C. upsaliensis* (n=3); *C. ureolyticus* (n=4); *C. volucris* (n=2); *Campylobacter sp* (n=1) (Supplementary File 1). Genomes belonging to *C. jejuni* and *C. coli* species were selected to represent a wide range of hosts, sequence types, and clonal complexes and reflect the known population structure for these two species. For other *Campylobacter* species, all genomes that were publicly available at the time of this study were included in the analysis. (Supplementary File 1).

### Pangenome characterization and phylogenetic analysis

Sequence data were analysed using PIRATE, a fast and scalable pangenomics tool which allows for orthologue gene clustering in divergent bacterial species [91]. Genomes were annotated in Prokka [92], using a genus database comprising well annotated *C. jejuni* strains NCTC11168, 81116, 81-176 and M1, and plasmids pTet and pVir in addition to the already existing databases used by Prokka [92]. Briefly, annotated genomes were used as input for PIRATE. Non-redundant representative sequences were produced using CD-HIT and the longest sequence was used as a reference for sequence similarity interrogation using BLAST/DIAMOND. Gene orthologues were defined as “gene families” and were clustered in different MCL thresholds, from 10 to 98 % sequence identity (10, 20, 30, 40, 50, 60, 70, 80, 90, 95, 98). Higher MCL thresholds were used to identify allelic variation within different loci. An inflation value of 4 was used to increase the granularity of MCL clustering between gene families. BLAST high-scoring pairs with a reciprocal minimum length of 90% of the query/subject sequence were excluded from MCL clustering to reduce the number of spurious associations between distantly related or conserved genes [93]. This information was used to generate gene presence/absence and allelic variation matrices. A core gene-by-gene multiple sequence alignment [89] was generated using MAFFT [94] comprising genes shared >95% of isolates. Phylogenetic trees, based on core gene-by-gene alignments, were reconstructed using the maximum-likelihood algorithm implemented in RAxML v8.2.11 [95] with GTRGAMMA as substitution model.

### Quantifying core and accessory genome variation

The degree of genetic differentiation between species was investigated gene-by-gene as in previous studies [40, 96] by calculating the average nucleotide identity (ANI) of all 631 *Campylobacter* genomes using FastANI v.1.0 [59]. The analysis generated a lower triangular matrix with the lowest ANI value at 75% (as computed by FastANI). A comparable gene presence/absence matrix was produced using PIRATE and was further used to generate a heatmap of accessory genome similarity based upon gene presence or absence. Subsequently, all *Campylobacter* genomes were screened for the presence of antimicrobial resistance genes against the CARD [62], ResFinder [63] and NCBI databases. All *Campylobacter* genomes were further screened for the presence of phage, conjugative elements and plasmid DNA using publicly available online databases to investigate the effect of other transfer mechanisms. First, we used the PHAge Search Tool Enhanced Release (PHASTER) [97] to identify and annotate prophage sequences within our genomes. A total of 86% (254/297) of the genomes used in chromosome painting were found to have DNA sequence of phage origin. Second, we used Iceberg 2.0 [98] for the detection of integrative and conjugative elements, identifying 32 ICEs in 19% (56/297) of the genomes used in the chromosome painting analysis. Finally, we used MOB-suite software for clustering, reconstruction and typing of plasmids from draft assemblies [99, 100]. A positive hit was defined when a gene had >75% nucleotide identity over >50% of the sequence length showing that 32 genes identified in the recombination analysis have also been located on plasmids. A gene presence/absence matrix for every antimicrobial resistance gene was generated for every genome. Genomes carrying AMR genes were screened to characterize the location of adjacent genes using SnapGene software (GSL Biotech; available at snapgene.com), as previously described [64]. The number of core SNPs was identified using SNP-sites (v2.3.2) [101].

### Inference of recombination

Each combination of a recipient group and multiple donor groups (belonging to the same CC but isolated from different hosts) was selected to compare the extent of interspecies recombination into the recipient genomes. Each donor group consisted of 8 isolates to avoid the influence of difference in sample size on estimation of the extent of interspecies recombination. Each recipient group included at least 4 isolates. We excluded *C. jejuni* and *C. coli* clade 1 genomes isolated from seals and water, as these most likely represent spillover events and not true host segregated populations. Briefly, we conducted a pairwise genome alignment between reference genome NCTC11168 and one of the strains included in the donor-recipient analysis using progressiveMauve [102]. This enabled the construction of positional homology alignments for all genomes regardless gene content and genome rearrangements, which were then combined into a multiple whole-genome alignment, as previously described [103]. ChromoPainterV2 software was used to calculate the amount of DNA sequence that is donated from a donor to a recipient group [45]. Briefly, for each donor-recipient pair, SNPs in which >90% recipient individuals had recombined with the donor group were considered in the analysis. These SNPs were mapped to genomic regions and specific genes were identified. A total of 258,444 (96.83%) recombining SNPs mapped to 558 genes of the NCTC11168 reference strain with >90% probability of copying from a donor to a recipient strain. Genes containing the highest number of recombining SNPs were considered for subsequent analyses (>95^th^ percentile) (Supplementary File 2). ClonalFrameML[104] was used to infer the relative number of substitutions introduced by recombination (*r*) and mutation (*m*) as the ratio *r*/*m* as previously described [16].

## Data availability

Genomes sequenced as part of other studies are archived on the Short Read Archive associated with BioProject accessions: PRJNA176480, PRJNA177352, PRJNA342755, PRJNA345429, PRJNA312235, PRJNA415188, PRJNA524300, PRJNA528879, PRJNA529798, PRJNA575343, PRJNA524315 and PRJNA689604. Additional genomes were also downloaded from NCBI [105] and pubMLST (http://pubmlst.org/campylobacter). Contiguous assemblies of all genome sequences compared are available at the public data repository Figshare (doi: 10.6084/m9.figshare.15061017) and individual project and accession numbers can be found in Supplementary File 1.

## Acknowledgements

This work was supported by Wellcome Trust grants 088786/C/09/Z and Medical Research Council (MRC) grants MR/M501608/1 and MR/L015080/1 awarded to S.K.S. The computational calculations were performed at the Human Genome Center at the Institute of Medical Science (University of Tokyo) and at the National Institute of Genetics.

---

**Figure 2–figure supplement 1. Population structure of the *Campylobacteraceae* family.** Phylogenetic tree of 506 isolates that belong to the *Campylobacteraceae* family with *Helicobacter pylori* used as an outgroup. Different colors correspond to main species with number of isolates greater than three. The Tree was reconstructed using a gene-by-gene concatenated alignment of 799 core genes shared by >95% by all isolates and an approximation of the maximum-likelihood algorithm (ML) implemented in RAxML. The scale bar indicates the estimated number of substitutions per site.

**Figure 2–figure supplement 2.** Core genome species trees. Single-species trees for nine *Campylobacter* species with >4 isolates demonstrating the diversity for among species. The scale bars indicate the estimated number of substitutions per site. (*) The scale for the tree corresponding to *C. hepaticus* is 10 times smaller than the rest.

**Figure 2–figure supplement 3. Overview of host-associations of *Campylobacter* species.** Abundance and diversity of 631 *Campylobacter* isolates in each host and environment. Different colours correspond to main species with number of isolates ≥ three. The number of isolates is shown on the *y* axis while the various isolation sources on the *x* axis.

**Figure 2–figure supplement 4.** Core genome species trees. Single-species trees for *C. jejuni, C. coli* and *C. fetus* species which contain isolates from multiple hosts and countries. The scale bars indicate the estimated number of substitutions per site.

**Figure 3–figure supplement 1. Genome size variation of the *Campylobacter* genus.** The frequency distribution of the genome size of all *Campylobacter* genomes used in this study is shown as a histogram. The number of genomes is shown on the *y* axis while the genome size (in bp) on the *x* axis.

**Figure 3–figure supplement 2. Gene variation in the genus *Campylobacter*.** Overall distribution of the total number of genes per isolate for each *Campylobacter* species (where n≥3 isolates). The number of genes is shown as boxplots (min to max).

**Figure 3–figure supplement 3. Accessory gene function in all main *Campylobacter* species.** The different gene functions are depicted on the *y* axis, while the number of shared accessory genes on the *x* axis. Different colours corresponding to different *Campylobacter* species.

**Figure 3–figure supplement 4. Core genome allelic variation and the effect of recombination. a**, Number of SNPs per genome of the main *Campylobacter* species (where n≥3 isolates) in the core genome alignment. The horizontal line in each plot represents the mean value while the upper and lower lines the standard deviation. **b**, Average nucleotide identity for pairwise comparisons of 820 core genes for 605 genomes of 15 main *Campylobacter* species. Different colours corresponding to different *Campylobacter* species.

**Figure 4-figure supplement 1.** Probability of the recipient genomes sharing DNA with each donor groups is illustrated as box whiskers (white) for every donor-recipient comparison for all 10 pairs that supported our hypothesis. The analysis where the host data were randomized across all isolates is illustrated as box whiskers (red). The probability of copying DNA from a donor to a recipient genome is shown on the *y* axis. The midline in the box whiskers indicates the mean and the error bars the standard deviation.

**Figure 4-figure supplement 2. Genome position of genes containing recombining SNPs.** Genes and their corresponding number of recombining SNPs, inferred by Chromosome Painting analysis for all 10 species comparisons, and mapped to the NCTC11168 reference genome. Genes from within-host (red) and between-host (white) pair comparisons are shown for each comparison. Donors are isolates from chicken (triangle), cattle (square), wild bird (cross), pig (star), clinical (circle) and water (snowflake) samples. The dashed line indicates the 95^th^ percentile for every individual group comparison.

**Figure 4-figure supplement 3.** Genes ranked in ascending order of the number of recombining SNPs they contain as inferred by Chromosome Painting analysis for all ten species comparisons. Genes from within-host (red) and between-host (white) are shown for each comparison. Donors are isolates from chicken (triangle), cattle (square), wild bird (cross), pig (star), clinical (circle) and water (snowflake) samples.

**Figure 5-figure supplement 1.** Presence of antimicrobial resistance genes in the *Campylobacter* genus. The phylogenetic tree was reconstructed using a gene-by-gene concatenated alignment of 820 core and soft-core genes and an approximation of the maximum-likelihood algorithm (ML) implemented in RAxML. The designated colour scheme was used for each species in the first column. The second column indicates whether the strain is isolated from an agricultural animal (grey). Remaining columns indicate presence of AMR genes (black). The scale represents the number of substitutions per site.

**Figure 5-figure supplement 2. Genetic organization of AMR genes in *Campylobacter*.** The presence of each AMR gene, highlighted in different colours, is shown for representative genomes from *C. jejuni, C. coli, C. lanienae, C. hyointestinalis* and *C. fetus subspecies fetus* sampled from different agricultural animals. The number of isolate genomes containing each genomic arrangement is indicated in parenthesis.

**Supplementary File 1. Isolate information about the genomes used in this study.**

**Supplementary File 2. Within-host highly (>95^th^ percentile) recombining genes.**

**Supplementary File 3. Recombination parameters as calculated by ClonalFrameML.**

**Supplementary File 4. Quantifying recombination between co-habiting species using ChromoPainter.**

**Supplementary File 5. Genes involved in interspecies recombination in 10 species comparisons.**

**Figure 3–source data 1. This file contains the numerical values on which the graphs in Figure 3 are based.**

**Figure 5–source data 1. This file contains the numerical values on which the graphs in Figure 5b-d are based.**

