## Supplementary figures and images for "Host ecology regulates interspecies recombination in bacteria of the genus *Campylobacter*"

### Figure 2-figure supplement 1

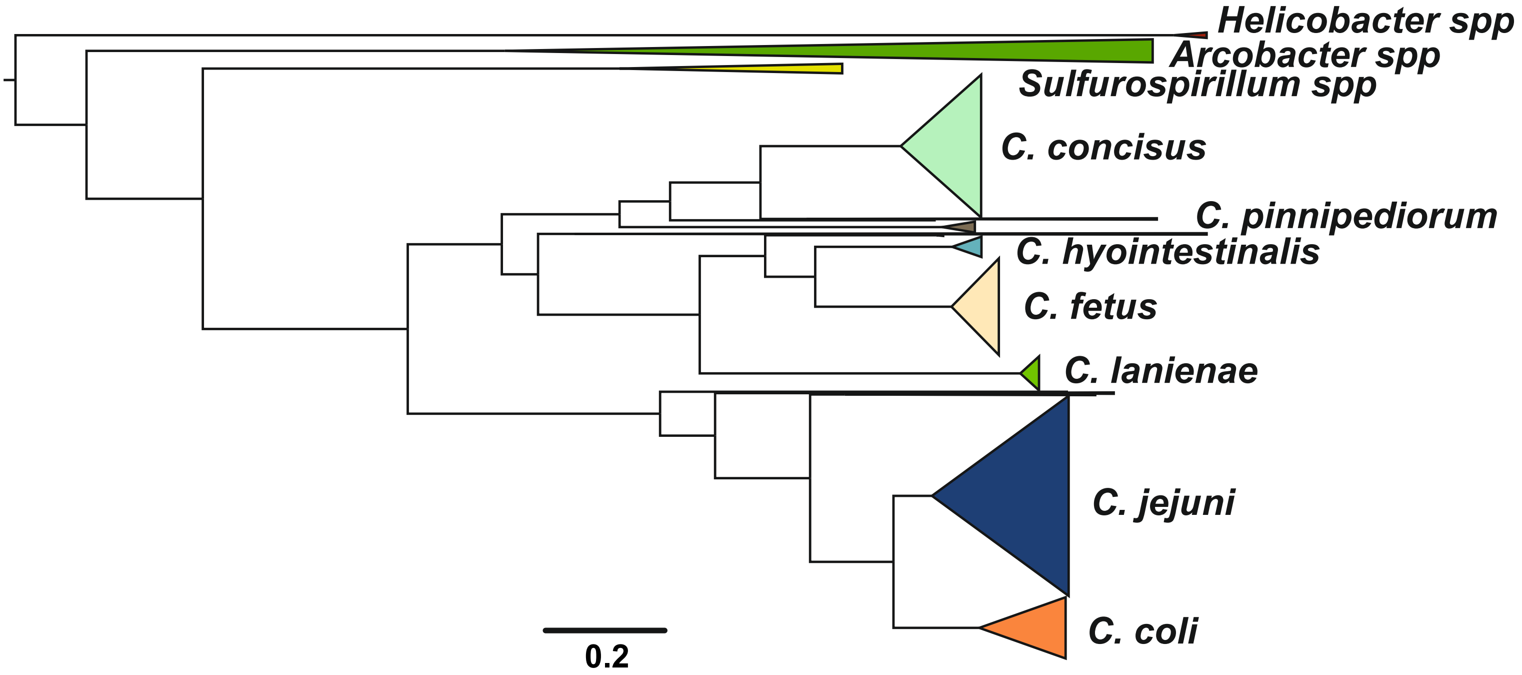

### Figure 2-figure supplement 2

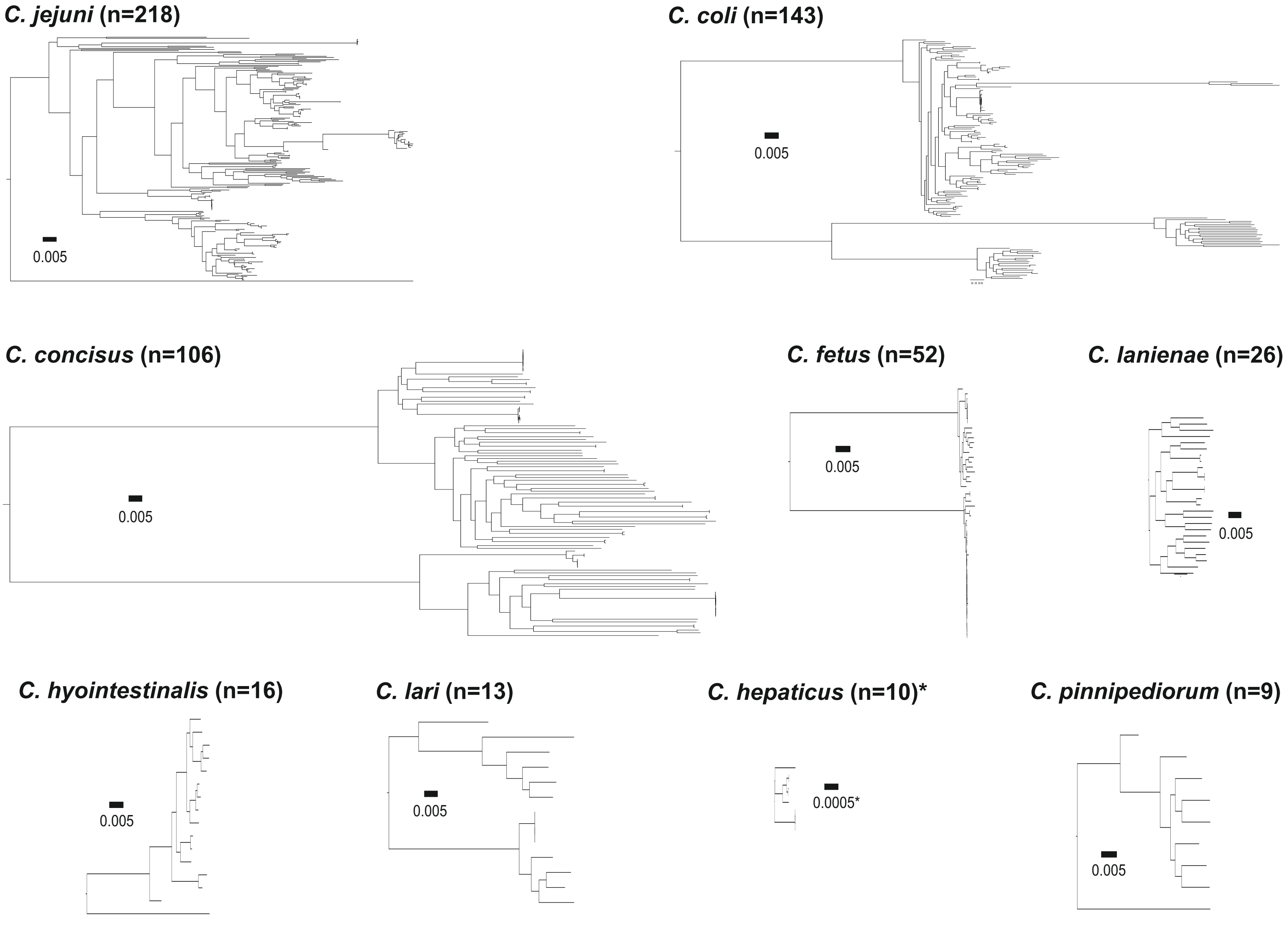

### Figure 2-figure supplement 3

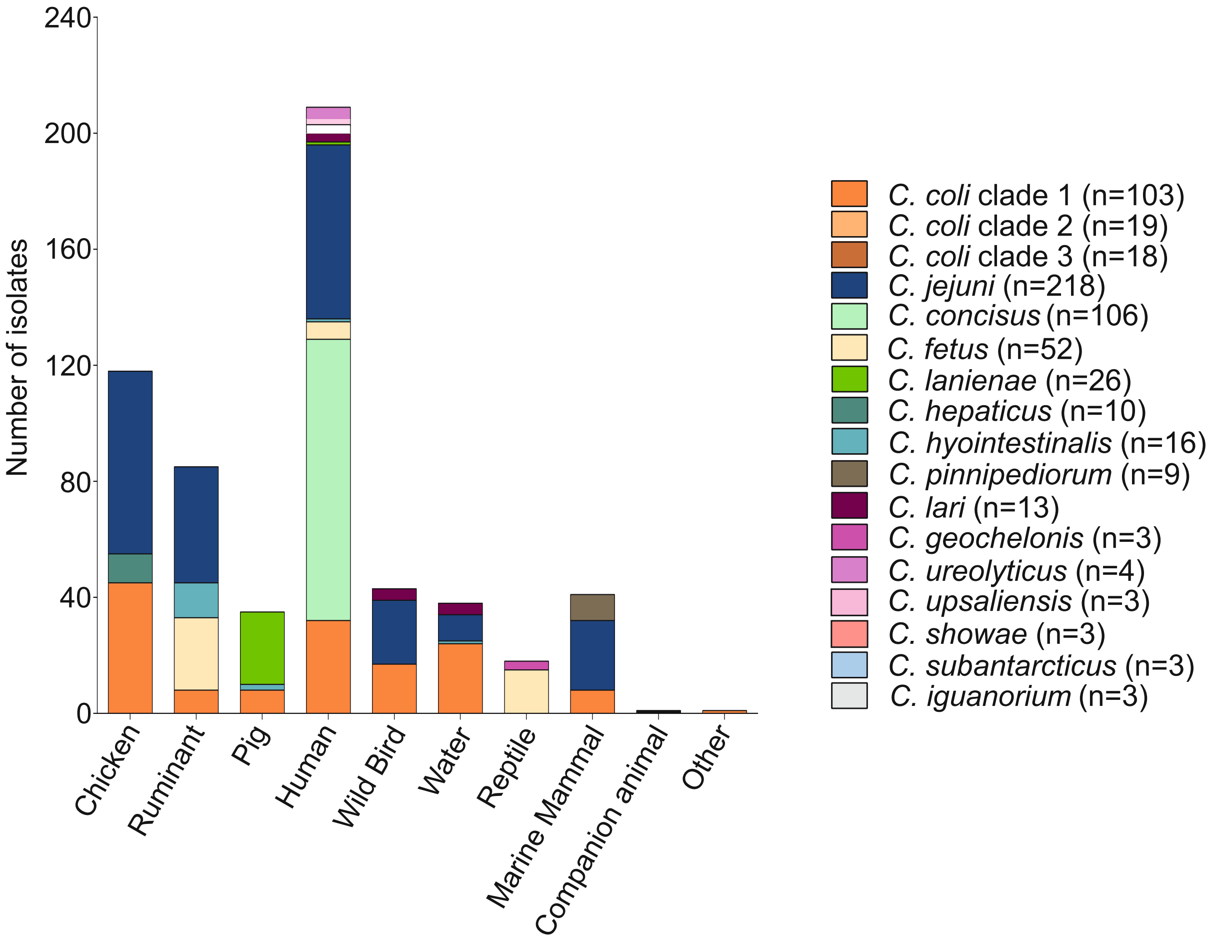

### Figure 2-figure supplement 4

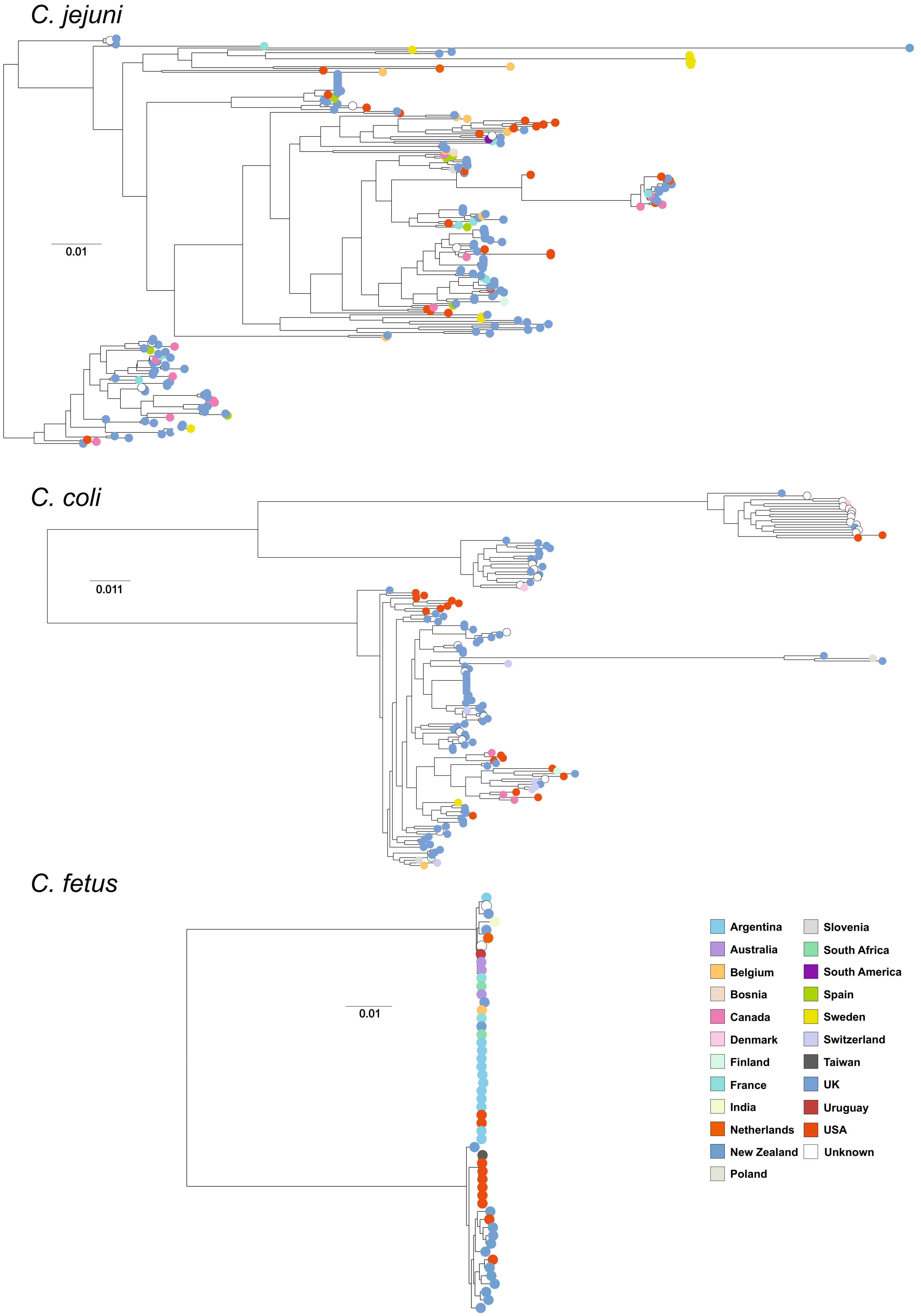

### Figure 3-figure supplement 1

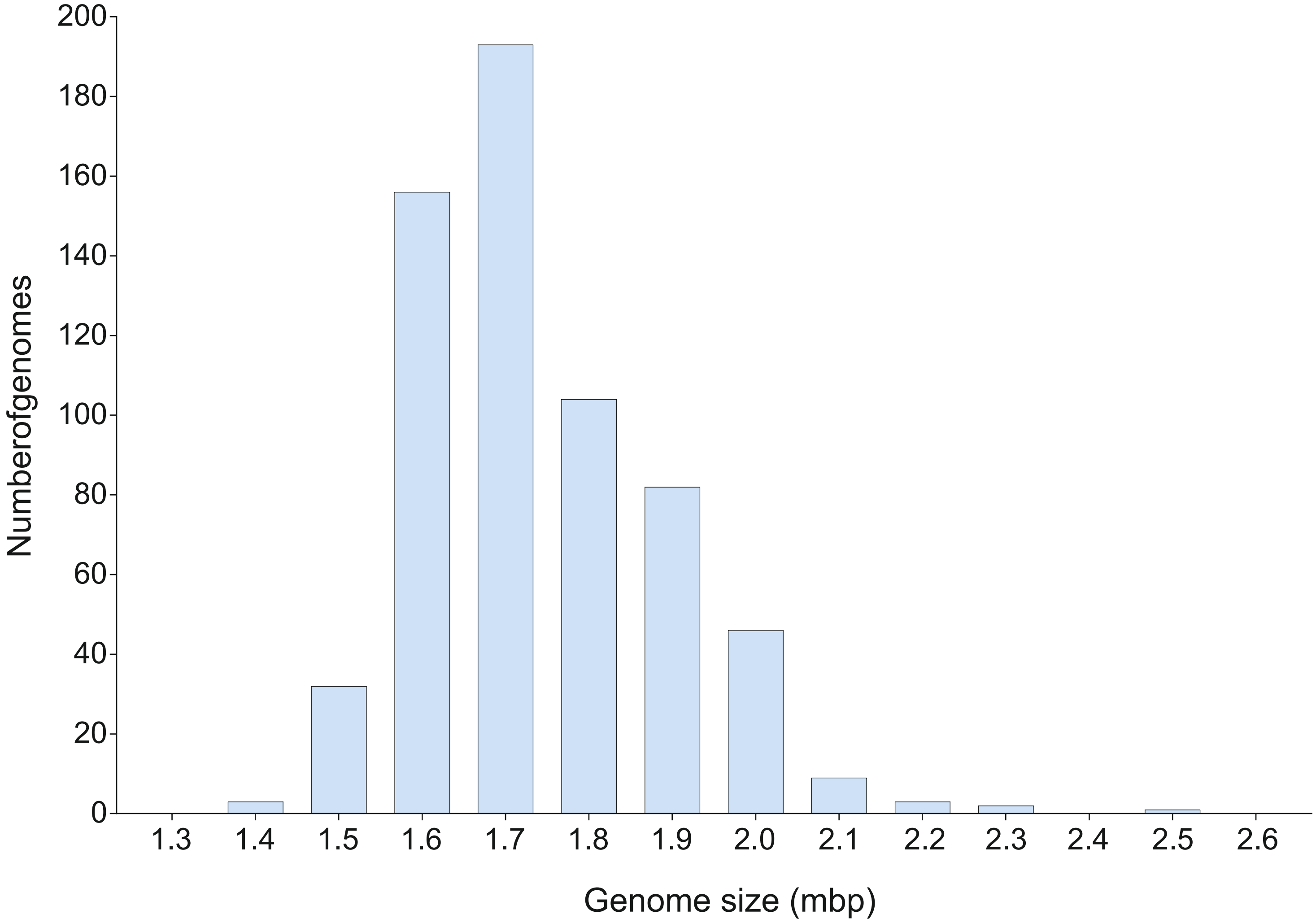

### Figure 3-figure supplement 2

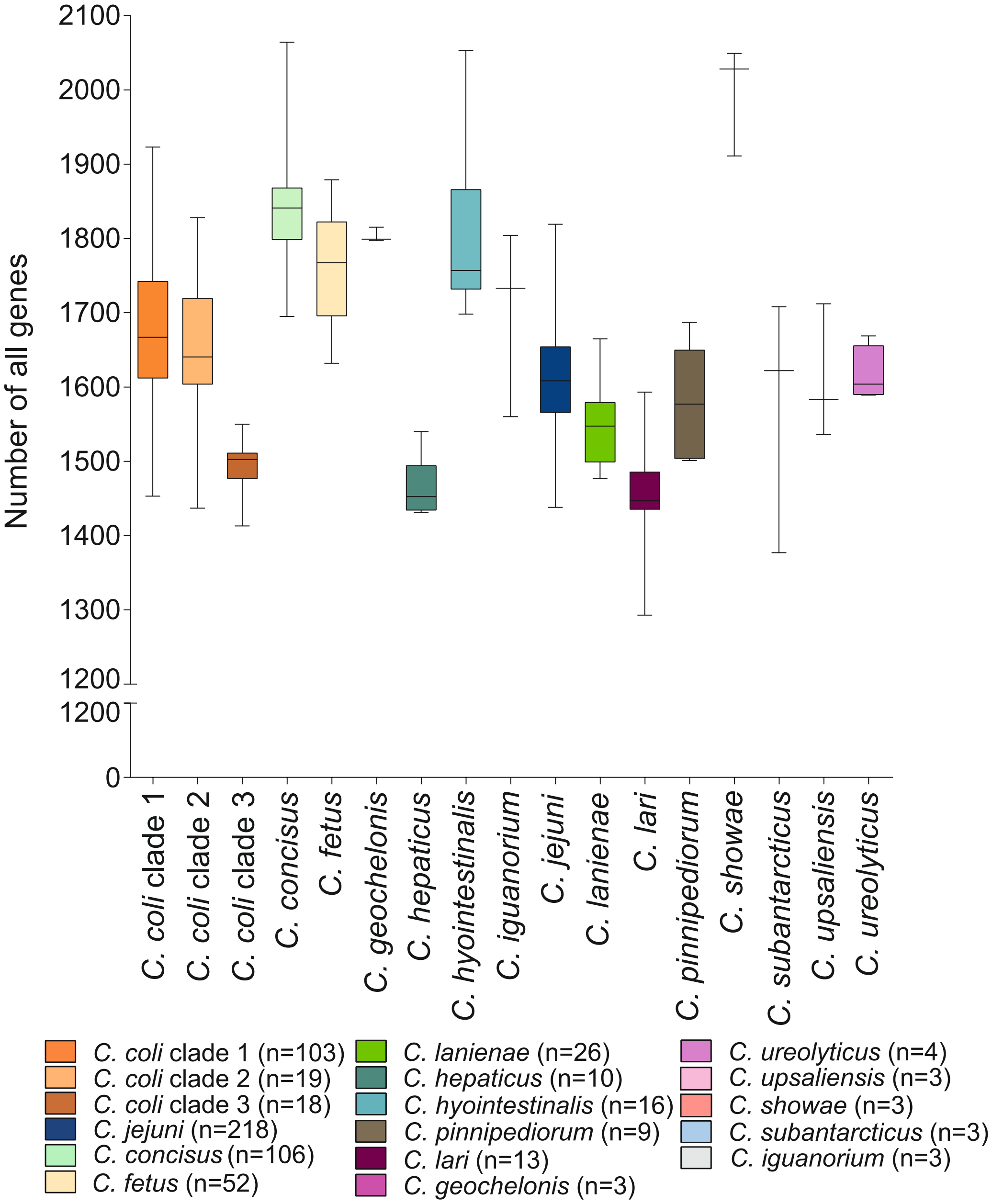

### Figure 3-figure supplement 3

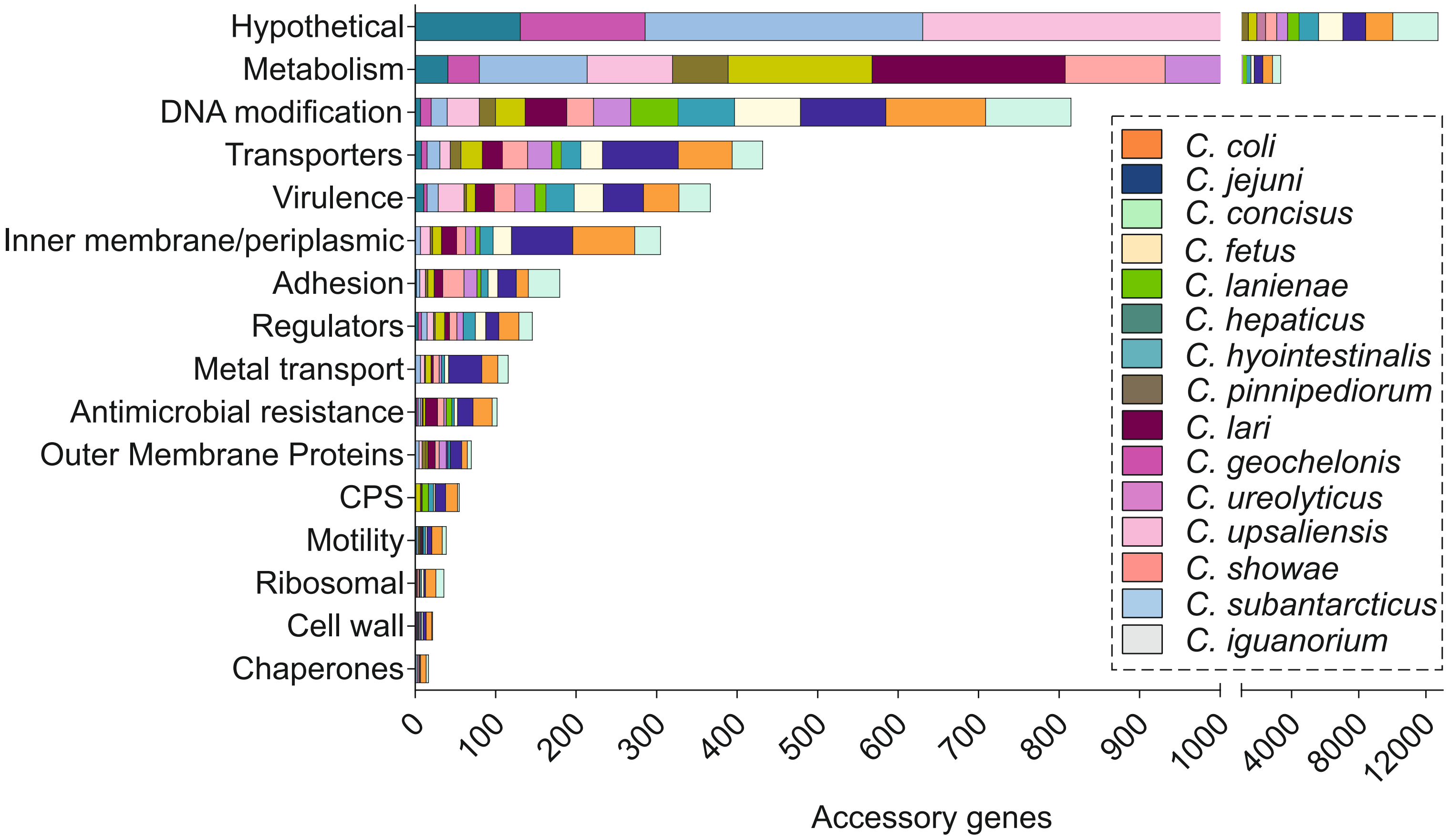

### Figure 3-figure supplement 4

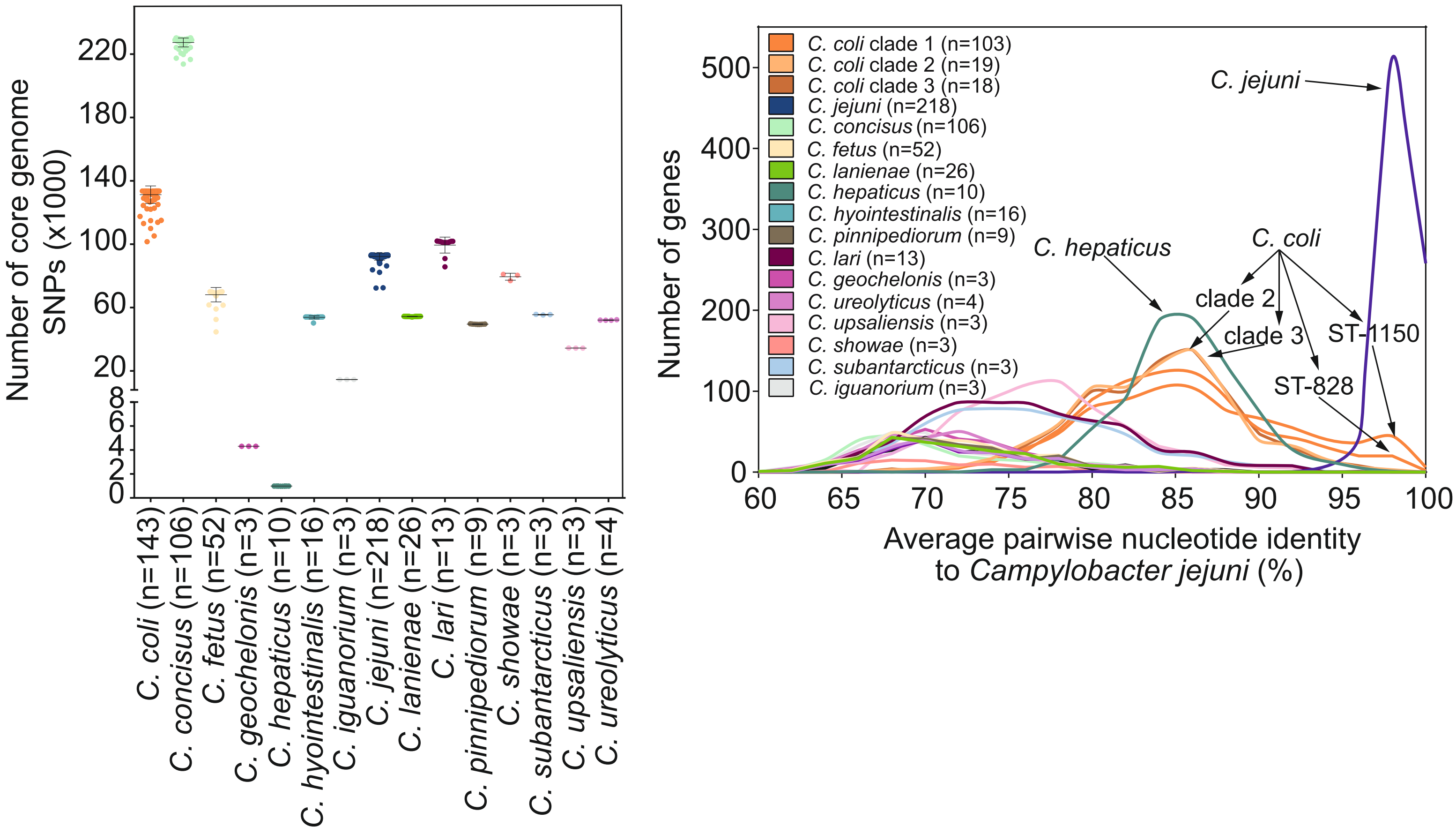

### Figure 4-figure supplement 1

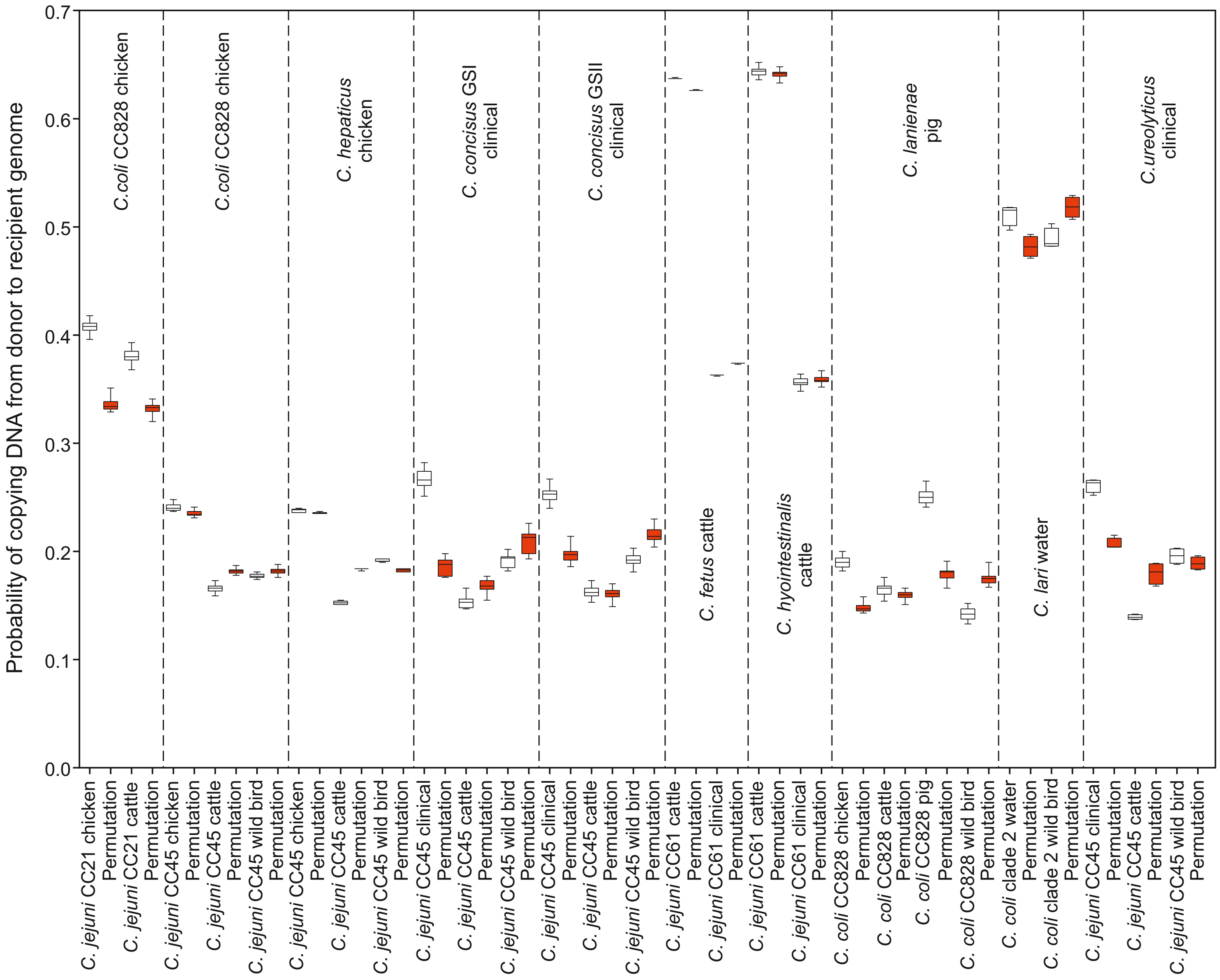

### Figure 4-figure supplement 2

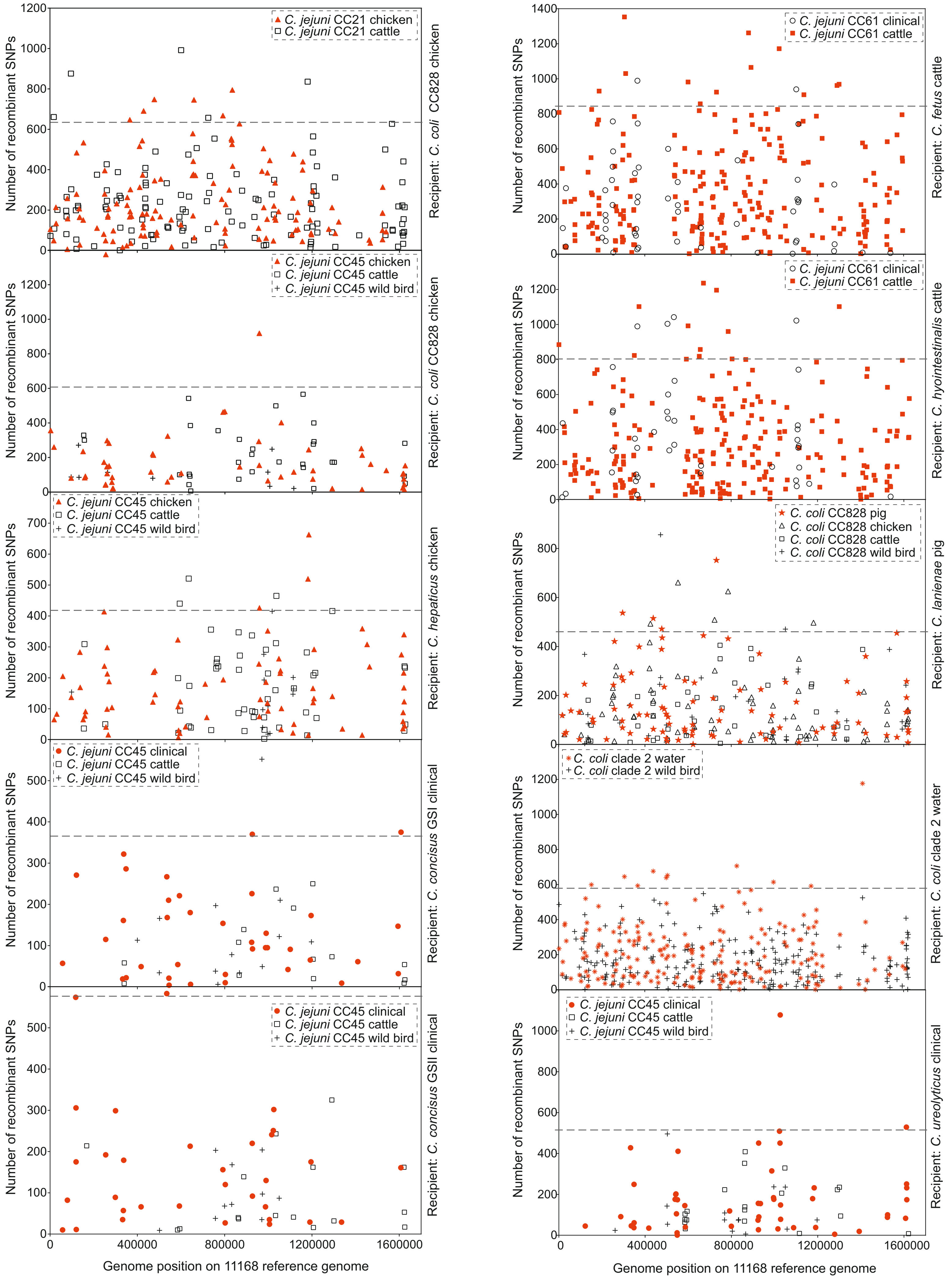

### Figure 4-figure supplement 3

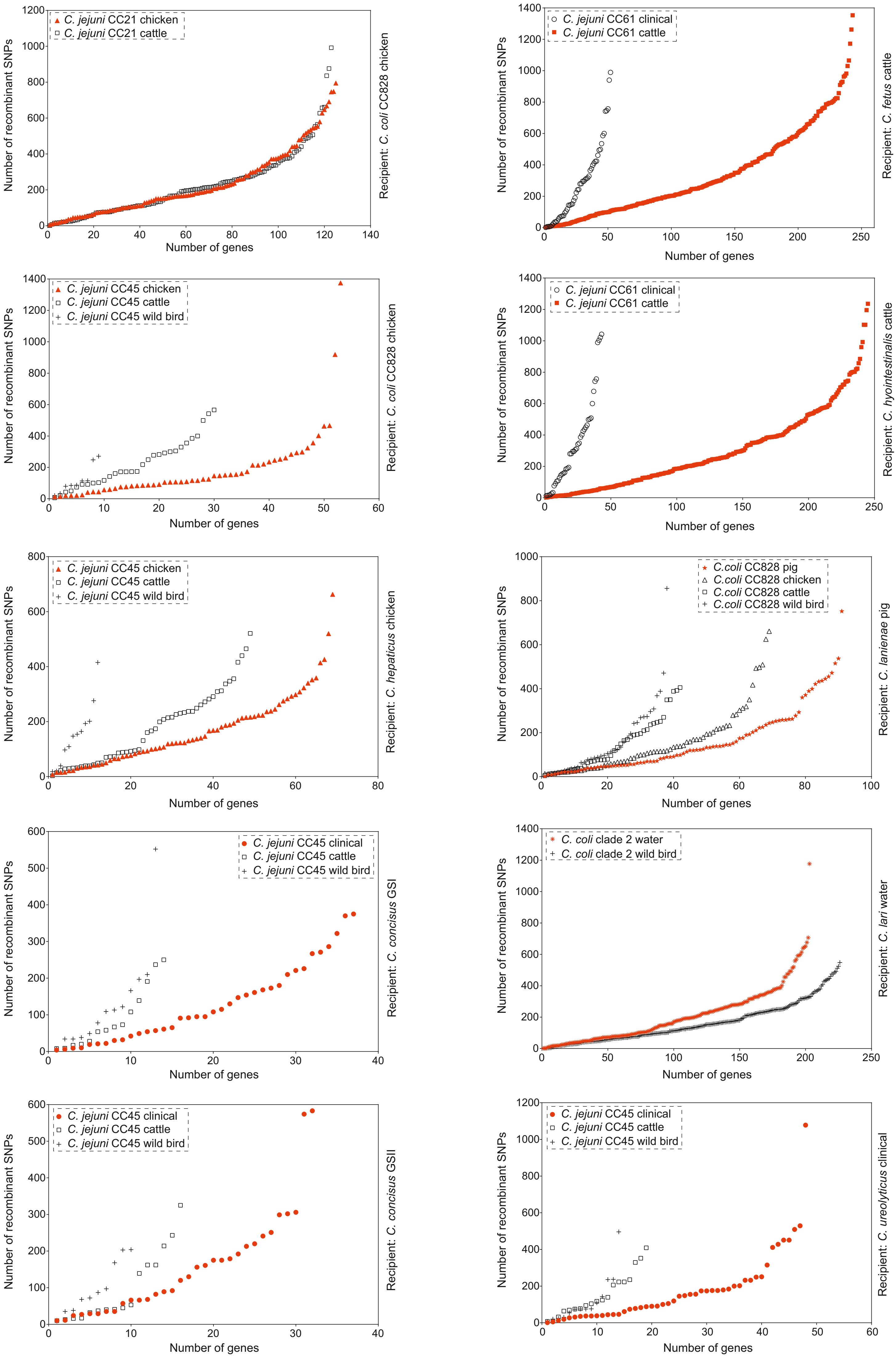

### Figure 5-figure supplement 1

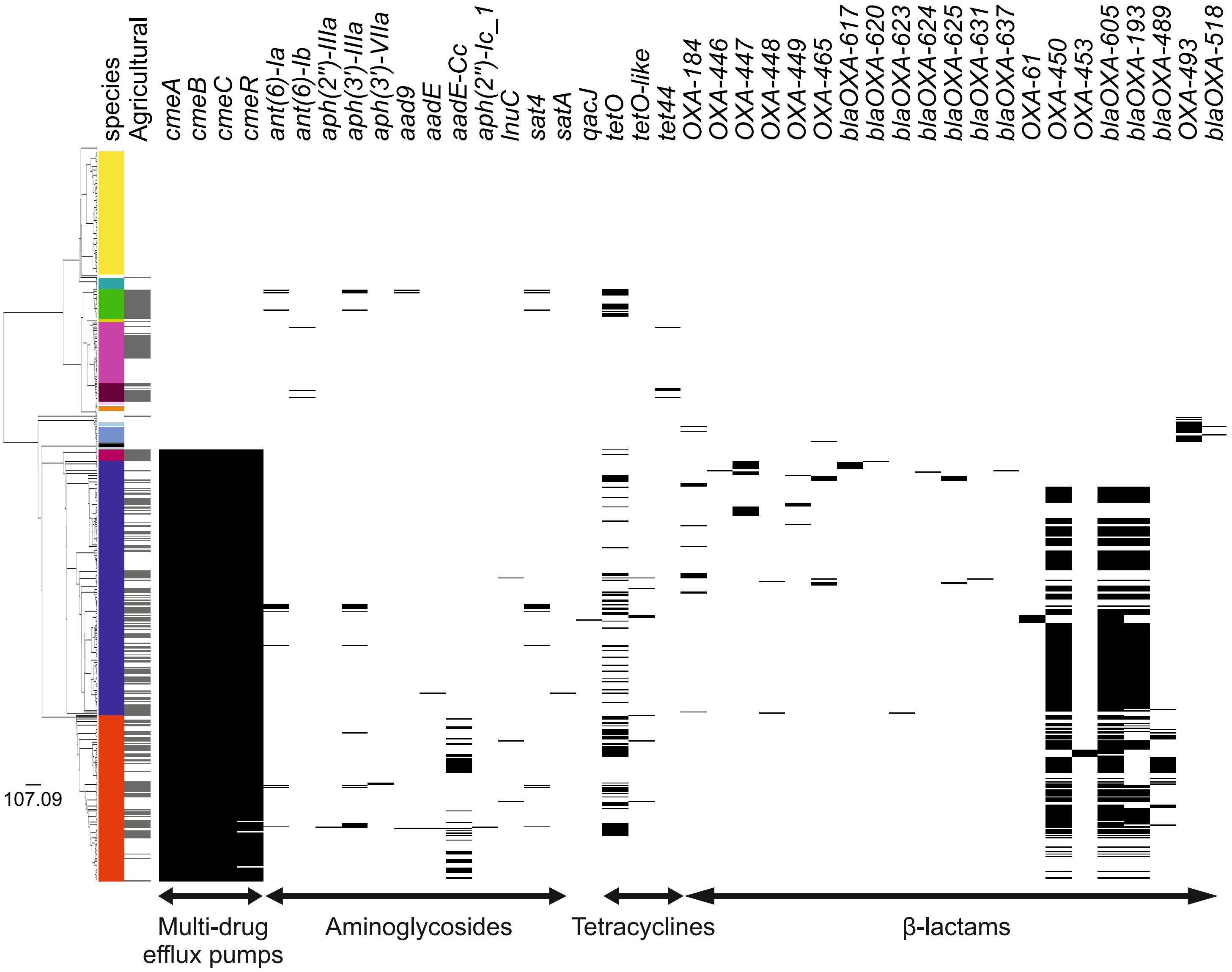

### Figure 5-figure supplement 2

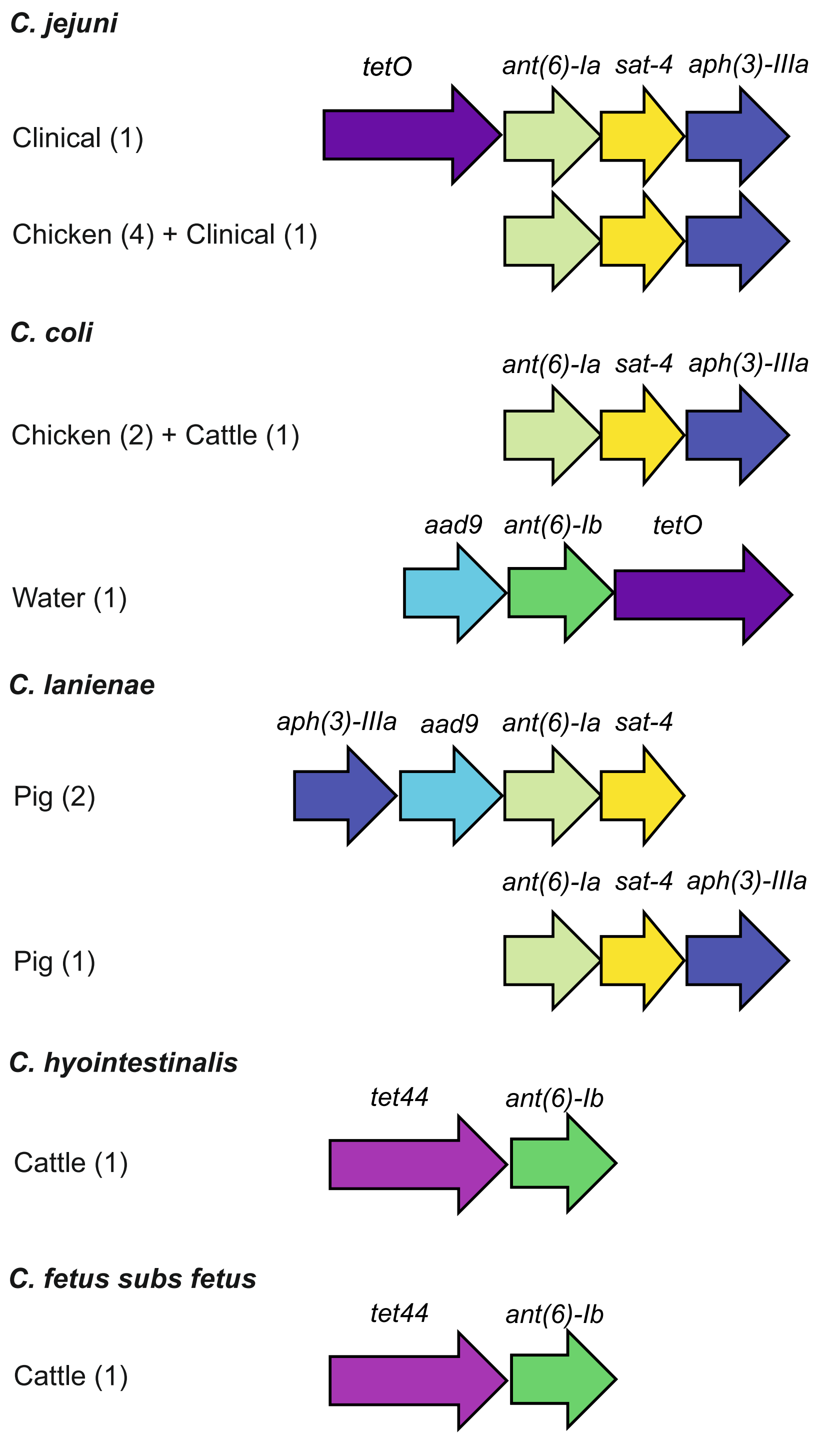
